# Knockdown of PHOX2B in the retrotrapezoid nucleus reduces the central CO_2_ chemoreflex in rats

**DOI:** 10.1101/2023.12.13.571591

**Authors:** Silvia Cardani, Tara A. Janes, William Betzner, Silvia Pagliardini

## Abstract

PHOX2B is a transcription factor essential for the development of different classes of neurons in the central and peripheral nervous system. Heterozygous mutations in the PHOX2B coding region are responsible for the occurrence of Congenital Central Hypoventilation Syndrome (CCHS), a rare neurological disorder characterised by inadequate chemosensitivity and life-threatening sleep- related hypoventilation. Animal studies suggest that chemoreflex defects are caused in part by the improper development or function of PHOX2B expressing neurons in the retrotrapezoid nucleus (RTN), a central hub for CO_2_ chemosensitivity.

Although the function of PHOX2B in rodents during development is well established, its role in the adult respiratory network remains unknown. In this study, we investigated whether reduction in PHOX2B expression in chemosensitive neuromedin-B (NMB) expressing neurons in the RTN altered respiratory function. Four weeks following local RTN injection of a lentiviral vector expressing the short hairpin RNA (shRNA) targeting *Phox2b* mRNA, a reduction of PHOX2B expression was observed in *Nmb* neurons compared to both naïve rats and rats injected with the non-target shRNA. PHOX2B knockdown did not affect breathing in room air or under hypoxia, but ventilation was significantly impaired during hypercapnia. PHOX2B knockdown did not alter *Nmb* expression but it was associated with reduced the expression of both *Task2* and *Gpr4*, two CO_2_ sensors in the RTN. We conclude that PHOX2B in the adult brain has an important role in CO_2_ chemoreception and reduced PHOX2B expression in CCHS beyond the developmental period may contribute to the impaired central chemoreflex function.

## INTRODUCTION

The Paired Like Homeobox 2B (PHOX2B) is a transcription factor essential for embryonic differentiation and for the maintenance of the neuronal phenotype of different classes of neurons in both the central and peripheral nervous systems (Brunet & Pattyn, 2002; Pattyn et al., 1997, 1999, 2000; Stanke et al., 1999; Tiveron et al., 1996). In recent years, the PHOX2B gene has been extensively investigated, not only for its neurodevelopmental role, but also because its heterozygous mutation leads to Congenital Central Hypoventilation Syndrome (CCHS, OMIM 209880; Amiel et al., 2003), Hirschsprung’s disease (HSCR; OMIM 142623; Berry-Kravis et al., 2006; Trochet et al., 2005), and neuroblastoma (Bourdeaut et al., 2005; Van Limpt et al., 2004). CCHS is a rare (1:200,000 births in France (Trang et al., 2005) and 1:148,000 births in Japan (Shimokaze et al., 2015) disorder characterised by a failure to respond to both hypercapnia and hypoxia (Amiel et al., 2003; Di Lascio et al., 2021; Weese-Mayer et al., 2017). The most frequent PHOX2B mutation is an expansion of a polyalanine repeat, ranging from +5 to +13 residues on the C-terminal of PHOX2B, an area that regulates DNA binding affinity and protein solubility (Amiel et al., 2003; Di Lascio et al., 2016; Weese-Mayer et al., 2003). CCHS symptoms present with varying degrees of severity depending on the expansion of the mutation (Berry-Kravis et al., 2006; Di Lascio et al., 2018; Matera, 2004). In most mild cases, patients display normal ventilation during wakefulness, hypoventilation during sleep leading to hypercarbia and hypoxemia, as well as a lack of sensitivity and discomfort to these challenges. In the most severe of cases, hypoventilation may also be present while awake.

Respiratory disturbances resulting from the PHOX2B mutations appear to stem from its widespread expression in respiratory-related neurons both during development and into adulthood. Although PHOX2B is absent in respiratory rhythmogenic neurons of the preBötzinger complex, it is highly expressed in adult neurons that are responsible for CO_2_ and O_2_ chemoreflexes (Kang et al., 2007). Among those are the neurons of the retrotrapezoid nucleus (RTN), the locus coeruleus, the carotid bodies and the nucleus of the solitary tract that, together with the dorsal motor nucleus of the vagus nerve and the area postrema, make up to the dorsal vagal complex, a structure that provides the major integrative centre for the mammalian autonomic nervous system (Cutsforth-Gregory & Benarroch, 2017; Guyenet et al., 2019; Kang et al., 2007; Zoccal et al., 2014). Previous work by our group and others specifically investigated the role of PHOX2B- expressing RTN neurons by either selective lesions or inactivation of neuronal activity of these neurons (Abbott et al., 2011; Basting et al., 2015; Marina et al., 2010; Nattie & Li, 1994; Souza et al., 2018, 2023; Janes et al., 2024) and showed that RTN neurons have a key role in central CO_2_ chemoreception.

Defects observed in the CO_2_-chemoreflex of CCHS transgenic rodent models are primarily caused by the improper development or function of PHOX2B-expressing neurons in the RTN (Dubreuil et al., 2008; Ramanantsoa et al., 2011), although anatomo-pathological studies in CCHS patients suggest that the respiratory impairment may be caused by a more widespread brain dysfunction (Harper et al., 2015; Nobuta et al., 2015). Transgenic mice born with PHOX2B knockout fail to develop the RTN, the locus coeruleus (Pattyn et al., 1999) and catecholaminergic groups in A1, A2, A5 and A7 (Pattyn et al., 2000), as well as neurons of the nucleus of the solitary tract and the carotid bodies. Mice harbouring heterozygous PHOX2B knockout develop apparently normal but have depressed responses to hypoxia and hypercapnia (Dauger, 2003) and sleep disordered breathing in the perinatal period (Durand et al., 2005). Furthermore, the expression of the most common heterozygous PHOX2B mutation found in CCHS patients (PHOX2B +7Ala expansion) in mice results in a near complete loss of RTN neurons, impaired CO_2_ response and consequent death in the newborn period (Dubreuil et al., 2008; Madani et al., 2021; Ramanantsoa et al., 2011; Takakura et al., 2008). More selective expression of the mutant PHOX2B in the RTN neurons allowed for survival in the neonatal period and partial recovery of the CO_2_ response in adulthood (Ramanantsoa et al., 2011).

Multiple potential cellular mechanisms have been implicated in the aetiology of CCHS, from loss of PHOX2B function in development due to heterozygous mutant PHOX2B expression, to gain of function of the mutated protein that may alter transcriptional activity in neurons, as well as aggregate formation and premature cell death (Bachetti et al., 2005; Di Lascio et al., 2013; Dubreuil et al., 2008; Durand et al., 2005; Goridis et al., 2010; Trochet et al., 2005). While its role during embryonic neurodevelopment is established, PHOX2B is still expressed in selected neuronal populations in the adult brain (Kang et al., 2007; Stornetta et al., 2006) and its function, with the exception of the contribution to the maintenance of the noradrenergic phenotype (Fan et al., 2011), is not yet fully understood. Considering the fundamental role of PHOX2B in CCHS pathogenesis and its expression in key structures for CO_2_ chemoreflex responses, it is essential to understand how both wild-type and mutant PHOX2B proteins impact CO_2_ homeostasis and ventilation not only across development, but also in adulthood.

In order to have a better understanding of the role of PHOX2B in the CO_2_ homeostatic processes we used a non-replicating lentivirus vector of two short-hairpin RNA (shRNA) clones targeting selectively *Phox2b* mRNA to knockdown the expression of PHOX2B in the RTN of adult rats and tested ventilation and chemoreflex responses. In parallel, we also determined whether knockdown of PHOX2B in adult *Nmb* neurons of the RTN negatively affected their cell survival. Finally, we sought to provide a mechanistic link between PHOX2B expression and the chemosensitive properties of *Nmb* neurons of the RTN, which have been attributed to two proton sensors, the proton-activated G protein-coupled receptor (GPR4) and the proton-modulated potassium channel (TASK-2) (Gestreau et al., 2010; Guyenet et al., 2016; Kumar et al., 2015). The expression of these sensors is partially overlapping in *Nmb* neurons of the RTN and the genetic deletion of either one impairs the central respiratory chemoreflex with maximal impairment when both sensors are eliminated (Gestreau et al., 2010; Guyenet et al., 2016; Kumar et al., 2015).

Our results indicate that progressive knockdown of PHOX2B in RTN causes a reduction in the chemoreflex response that was proportional to the decrease in the fraction of *Nmb*/PHOX2B expressing RTN neurons. This effect was associated with a reduction in the expression of *Gpr4* and *Task2* mRNA in *Nmb* expressing cells of the RTN in which PHOX2B was knocked down, suggesting a role for PHOX2B in the transcription of the proton sensors in RTN neurons.

## RESULTS

To directly investigate the role of PHOX2B in RTN neurons, we performed selective knockdown of this protein in adult rats through local injection of a non-replicating lentiviral vector of two short- hairpin RNA (shRNA) clones targeting two different sequences of the *Phox2b* mRNA (n=25) or non- target shRNA as control (NT-shRNA; n=23). Initial experiments utilized 200 nl/injection at each of 4 stereotaxic coordinates (1x10^9^ VP/ml; n=14). Four weeks post injection, we assessed ventilatory parameters in room air, hypercapnia, and hypoxia. Interestingly, a small but significant impairment in the CO_2_ chemoreflex was observed not only in PHOX2B-shRNA but also in NT-shRNA rats compared to pre-surgical baseline (allometric V_E_ in 7% CO_2_: -21% and -24%, respectively; Supp. Fig. 1A). Because histological analysis also showed significant reduction of *Nmb*^+^/PHOX2B^+^ RTN cells in NT-shRNA and PHOX2B-shRNA rats (33% and 19% cells remaining, respectively; Suppl. Fig. 1), we concluded that the volume and titre of the shRNA viral constructs triggered off-target effects (Mcbride et al., 2008; Van Gestel et al., 2014) that cause cell death in RTN neurons.

### PHOX2B expression is modestly reduced two weeks following infection without respiratory impairment

Due to the off-target effects obtained with large volume injections, we reduced injection volume of both PHOX2B shRNA (n=17) and NT-shRNA viruses (n=17) (4x100nl/injection; 1x10^9^ VP/ml).

Two weeks post-surgery we assessed respiratory function during exposure to room air, hypercapnia and hypoxia. Overall, we observed little change in respiration at this time point. During room air, as well as in 5% and 7.2% CO_2_, frequency (ƒ_R_), was reduced in every experimental group (naïve, NT-shRNA, PHOX2B-shRNA) compared to baseline (Air: p=0.014; 5% CO_2_: p< 0.001; 7.2% CO_2_ p< 0.001; Fig. 1A) but no effect on treatment was observed, suggesting that these changes were likely due to the increase in age and body weight during the long-term experiment (naïve: +32.2%; NT-shRNA: +27.8%; PHOX2B-shRNA: +28.6% change in body weight). Tidal volume was reduced in the PHOX2B-shRNA rats at 7.2% CO_2_ compared to both naïve rats (naive: 9.6 ± 1.2 ml·kg^-1^; PHOX2B-shRNA: 8.5 ± 0.9 ml·kg^-1^; -11%; p= 0.022) and baseline pre-surgery conditions (baseline: 9.6 ± 1.3 ml·kg^-1^; -11%; p< 0.001; Fig. 1B) but it was not different from NT-shRNA.

**Figure 1.**
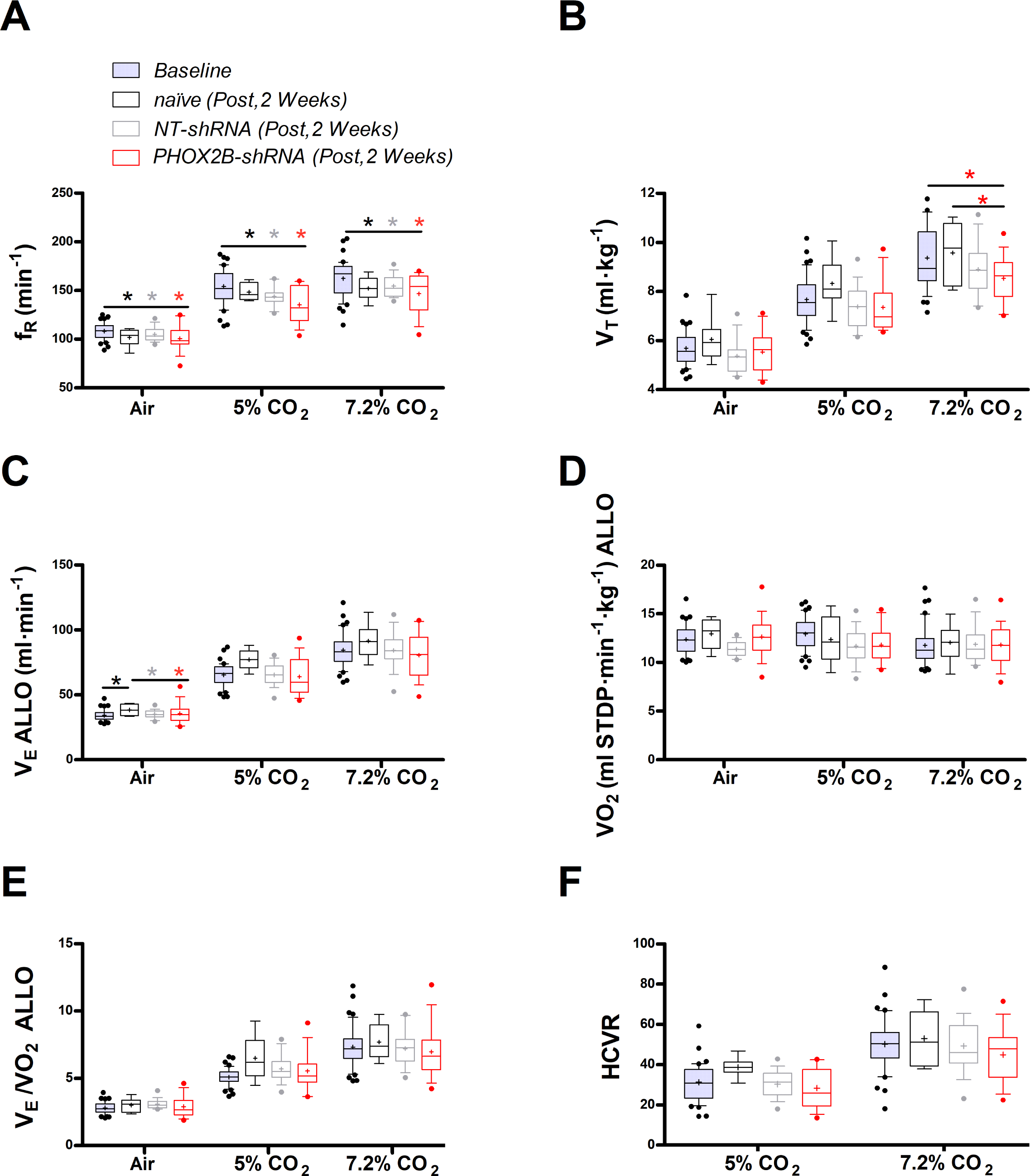
**Respiratory data two weeks post viral PHOX2B shRNA injection**. **(A)** breathing frequency (ƒ_R_,), **(B)** tidal volume (V_T_), **(C)** allometric minute ventilation (V_E_ ALLO), **(D)** oxygen consumption (VO_2_ ALLO), **(E)** convective requirement ratio (V_E_/VO_2_ ALLO), **(F)** hypercapnic ventilatory response (HCVR absolute change in V_E_ ALLO vs. corresponding room air) of baseline (pre-surgery, grey filled box), naïve (black n=8), non-target control shRNA (NT-shRNA, grey n=17) and PHOX2B shRNA (PHOX2B-shRNA, red n=17) rats 2-weeks post injection during room air, hypercapnia 5% and 7.2% CO_2_. ƒ_R_ is equally impaired in all experimental group compared to baseline but no treatment effect was observed (A). V_T_ is significantly impaired following RTN injection in PHOX2B-shRNA group compared to naïve animals (p=0.022) and baseline (p< 0.001) at 7.2% CO_2_ (B). V_E_ ALLO is increased in naïve rats compared to the other treatment groups (p=0.005) in room air (C). Boxplots: median, 1^st^ – 3^rd^ quartiles and 10^th^ – 90^th^ percentiles, outliers = dots, “+” indicates arithmetic mean. Bonferroni post-hoc as indicated. Black*, different from naïve; Grey*, different from NT-shRNA; Red*, different from PHOX2B- shRNA.

Despite similar changes in body weight across treatment groups, allometric ventilation (V_E_ ALLO) was higher in naïve rats compared to the other treatment groups in room air (naïve: 38.4 ± 4.5 ml·min^-1^; NT-shRNA: 34.7 ± 2.8 ml·min^-1^; PHOX2B-shRNA: 34.0 ± 4.5 ml·min^-1^; p=0.005; Fig. 1C), but no changes in V_E_ ALLO were observed in hypercapnia (7.2% CO_2_) across treatments (naïve : 91.5 ± 13.2 ml·min^-1^; NT-shRNA: 84.4 ± 13.7 ml·min^-1^; PHOX2B-shRNA: 80.5 ± 17.2 ml·min^-1^). Oxygen consumption (VO_2_ ALLO) was not different between experimental conditions or treatment groups (Fig. 1D), and V_E_/VO_2_ ALLO indicated that ventilation was matched with metabolism for each treatment across all gas compositions (naïve: 7.7 ± 1.3; NT-shRNA: 7.2 ± 1.3; PHOX2B-shRNA: 6.9 ± 2.0; Fig. 1E). These results suggest that despite a reduction in hypercapnic V_T_ in the PHOX2B- shRNA rats, overall ventilation was not impaired. Since we observed differences in room air V_E_ ALLO between treatment groups, we normalized these data to show the hypercapnic ventilatory response (HCVR, i.e., absolute change in V_E_ from room air; Fig. 1F). The HCVR was also not affected by the PHOX2B shRNA treatment two weeks post-injection (naïve: 53.1 ± 13.4 ml·min^-1^; NT-shRNA: 49.6 ± 13.2 ml·min^-1^; PHOX2B-shRNA: 46.5 ± 14.8 ml·min^-1^).

To determine the extent of PHOX2B knockdown in RTN neurons we combined RNAScope^®^ and immunohistochemistry assays to quantify RTN neurons expressing *Nmb* and PHOX2B (Fig. 2B). We reasoned that by assessing the fraction of RTN neurons that express only *Nmb* (compared to *Nmb*^+^/PHOX2B^+^), we would be able to determine the level of PHOX2B knockdown in our experiments. Two weeks post viral injection, we observed a small decrease (15.2%) in the total number of *Nmb* RTN neurons (i.e., *Nmb*^+^/PHOX2B^+^ plus *Nmb*^+^/ PHOX2B^-^) in NT-shRNA rats compared to naïve rats (naïve; 337.6 ± 11.95; n=4; NT-shRNA: 286.0 ± 10.44; n=4; p<0.001; Fig. 2C). The total number of *Nmb* cells of the RTN in PHOX2B-shRNA rats was further reduced compared to both naïve rats (PHOX2B-ShRNA 218.8 ± 7.8 n=4; -35.2%; p<0.001) and to NT-shRNA (-23.5%; p<0.001).

**Figure 2.**
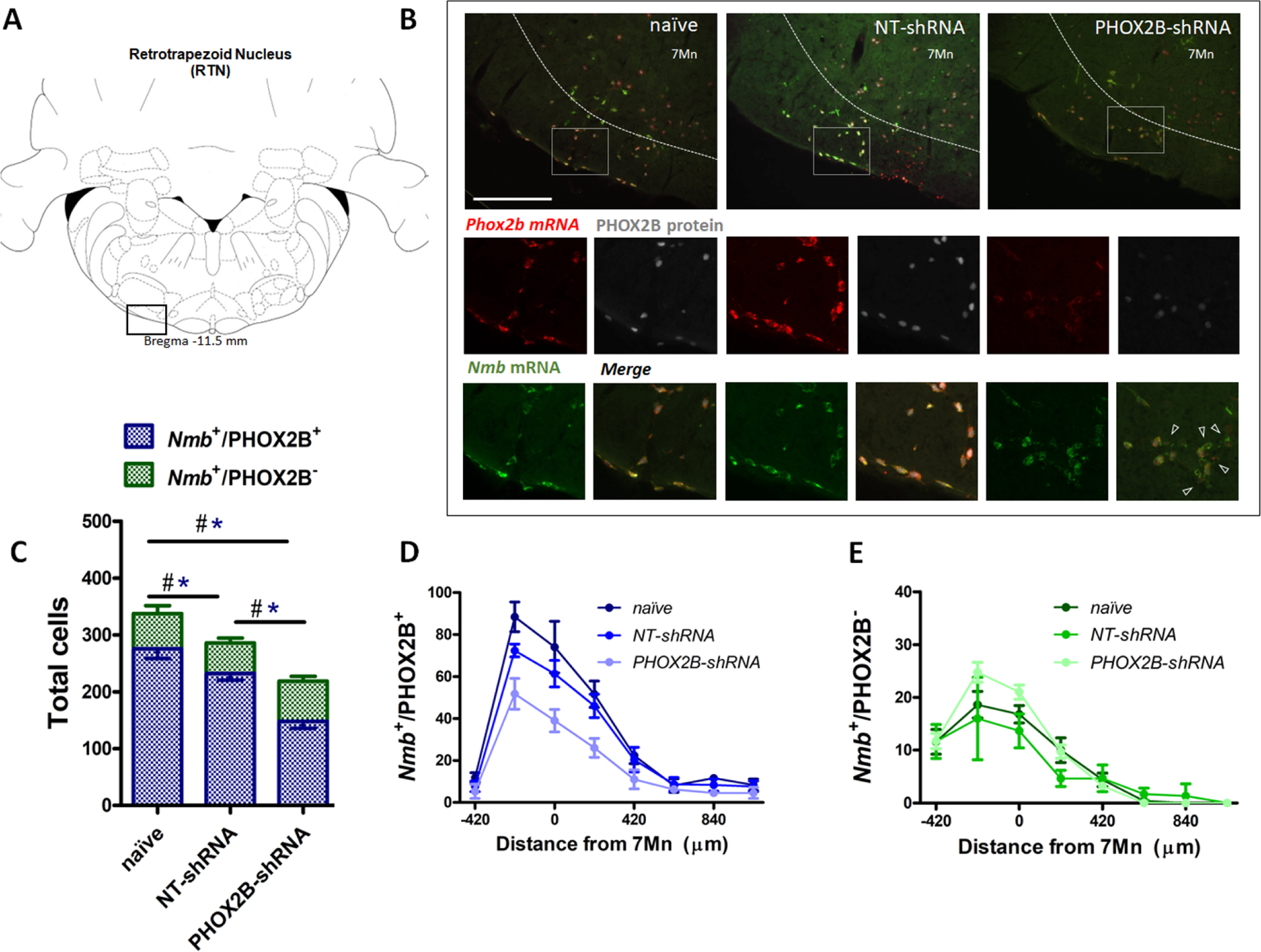
PHOX2B and *Nmb* expression and total cell count within the RTN area in naïve, NT- shRNA and PHOX2B-shRNA injected rats two weeks post viral shRNA injection. (A) Schematic and representative image of a transverse brainstem section at the level of the RTN (-11.5 mm distance from Bregma) showing the area of investigation containing RTN neurons. **(B)** Expression of *Phox2b* mRNA (red), PHOX2B protein (white) and *Nmb* mRNA (green) in RTN *Nmb*^+^/PHOX2B^+^ and *Nmb*^+^/PHOX2B^-^ neurons in naïve (left), NT-shRNA (middle), and PHOX2B-shRNA (right) rats (magnified view inserts below). Arrowheads indicate absence of PHOX2B protein. Scale bar=400μm (top figures), 150μm (inserts below). **(C)** The number of total cells (*Nmb*^+^/PHOX2B^+^ plus *Nmb*^+^/PHOX2B^-^) comprising the RTN are reduced in PHOX2B-shRNA rats (n=4) as compared to naïve (n=4) and NT-shRNA (n=4), and in NT-shRNA rats as compared to naïve rats (black^#^, One-way ANOVA, p<0.001). The number of *Nmb*^+^/PHOX2B^+^ cells are reduced in PHOX2B-shRNA rats as compared to naïve and NT-shRNA (Blue*, One-way ANOVA, p<0 0.001). The number of *Nmb*^+^/PHOX2B^-^ cells was unchanged across groups. **(D,E)** Rostral-caudal distribution (distance from the caudal tip of the facial nucleus, 7Mn) of *Nmb*^+^/PHOX2B^+^ (D) and *Nmb*^+^/PHOX2B^-^ (E) neurons along the RTN.

We then quantified the number of neurons in the RTN expressing either *Nmb* and PHOX2B (*Nmb*^+^/PHOX2B^+^) or *Nmb* only (*Nmb*^+^/PHOX2B^-^). The number of *Nmb*^+^/PHOX2B^+^ neurons in PHOX2B-shRNA rat was reduced by 36.1 % compared to NT-shRNA and by 46.2% compared to naïve rats (*Nmb*^+^/PHOX2B^+^ NT-shRNA: 232.3 ± 11.6 vs naïve: 275.8 ± 17.1; vs PHOX2B-shRNA: 148.3 ± 12.6; p<0.05 and 0.001, respectively; Fig. 2C,D). On the contrary, the fraction of *Nmb*^+^/PHOX2B^-^ neurons in PHOX2B-shRNA rat was not different compared to naïve and NT-shRNA rats (*Nmb*^+^/PHOX2B^-^ NT-shRNA: 53.7 +- 8.6; vs naïve: 61.8 +-13.9; vs PHOX2B-shRNA: 70.5 +- 8.7 cells; Fig. 2C,E). These results thus indicate that despite modest loss of *Nmb*^+^ neurons in NT-shRNA and a reduction in the proportion of *Nmb* neurons expressing PHOX2B in PHOX2B-shRNA rats, 2 weeks of PHOX2B shRNA viral infection was not sufficient to impair ventilation.

### Four weeks of PHOX2B knockdown impairs the CO_2_-chemoreflex

Since the shRNAs integrate into the genome of the infected cells and are continuously produced, it is reasonable to expect that knockdown efficiency is time-dependent, and an extended infection time would result in a higher knockdown effect. To test this hypothesis, a subgroup of rats (naïve, NT-shRNA and PHOX2B-shRNA) were investigated four weeks post-viral injection (Fig. 3A,B).

**Figure 3.**
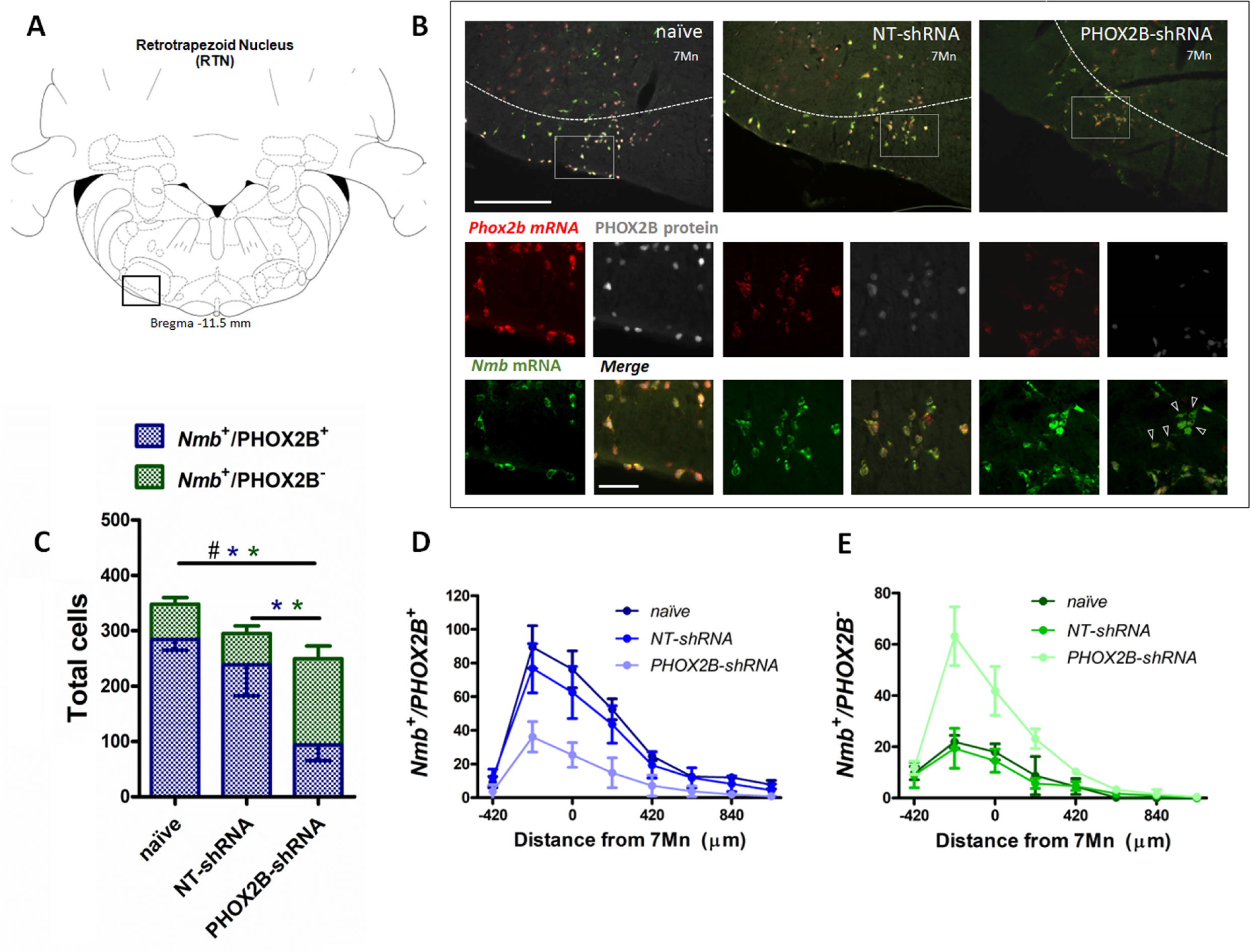
PHOX2B and *Nmb* expression and total cell count within the RTN area in naïve, NT- shRNA and PHOX2B-shRNA injected rats 4 weeks post viral shRNA injection. (A) Schematic and representative image of a transverse brainstem section at the level of the RTN (-11.5 mm distance from Bregma) showing the area of investigation containing RTN neurons. **(B)** Expression of *Phox2b* mRNA (red), PHOX2B protein (white) and *Nmb* mRNA (green) in naïve (left), NT-shRNA (middle), and PHOX2B-shRNA (right) rats (magnified view insert). Arrowheads indicate absence of PHOX2B protein. Scale bar = 400μm (top figures), 150μm (inserts below). **(C)** The number of total cells (*Nmb*^+^/PHOX2B^+^ + *Nmb*^+^/PHOX2B^-^) comprising the RTN are reduced in PHOX2B-shRNA rats (n=6) as compared to naïve (n=4) (Black^#^, One-way ANOVA, p=0.0087) but not to NT-shRNA (n=10). The number of *Nmb*^+^/PHOX2B^+^ cells are reduced in PHOX2B-shRNA rats as compared to naïve and NT- shRNA (Blue*, one-way ANOVA, p<0.001). The number of *Nmb*^+^/PHOX2B^-^ cells are increased in PHOX2B-shRNA rats as compared to both naïve and NT-shRNA (Green*, one-way ANOVA p<0.001). **(D, E)** Rostral-caudal distribution (distance from the caudal tip of the facial nucleus, 7Mn) of *Nmb*^+^/PHOX2B^+^ (D) and *Nmb*^+^/PHOX2B^-^ (E) neurons along the RTN.

The total number of *Nmb* neurons in the RTN of PHOX2B-shRNA rats (*Nmb*^+^/PHOX2B^+^ plus *Nmb*^+^/PHOX2B^-^; Fig. 3C) was significantly reduced compared to naïve rats (naïve: 347.8 ± 12 cells, n=4; PHOX2B-shRNA: 249.5 ± 33.9 cells; n=6; -28.3%; p=0.009) but no difference was observed compared to NT-shRNA rats (294.9 ± 52.8 cells; n=10; -15.4%;). Furthermore, there was no difference in the total number of *Nmb* cells in RTN between 2 and 4 weeks post infection for both NT-shRNA (week 2: 286 ± 10.4; week 4: 294.9 ± 52.8, p=0.783) and PHOX2B-shRNA rats (week 2: 218.8 ± 7.8; week 4: 249.5 ± 33,9, p=0.118), suggesting that the progressive PHOX2B knockdown was specific for PHOX2B-shRNA treatment and it was not accompanied by an increased cell death beyond the first 2 weeks.

The number of *Nmb*^+^/PHOX2B^+^ cells in the RTN was significantly reduced in PHOX2B-shRNA rats by 67.0% and 60.7% compared to naïve and NT-shRNA rats, respectively (naïve: 284.5 ± 19.8; NT- shRNA: 238.7 ± 56; PHOX2B-shRNA: 93.8± 11.9 cells, p<0.001; Fig. 3D). Moreover, a significant increase in the fraction of *Nmb*^+^/PHOX2B^-^ was observed with PHOX2B knockdown (PHOX2B- shRNA: 155.7 ± 22.7; naïve: 63.2 ± 12cells; + 146.1% increase vs naïve; NT-shRNA: 56.2 ± 13.8cells; + 177% increase vs NT-shRNA, p<0.001; Fig. 3E), confirming the efficiency of the PHOX2B knockdown in the *Nmb* cells of the RTN.

The respiratory function prior to histological examination (4 weeks post-viral injection) showed significant differences in PHOX2B-shRNA rats (n=6) compared to both pre-surgery baseline, naïve (n=8) and NT-shRNA rats (n=10). Similar to what we reported at 2 weeks post-surgery, ƒ_R_ at 4 weeks was lower in every treatment group compared to baseline recordings (Air: p=0.001; 5% CO_2_: p< 0.001; 7.2% CO_2_ p< 0.001; Fig. 4A).

**Figure 4.**
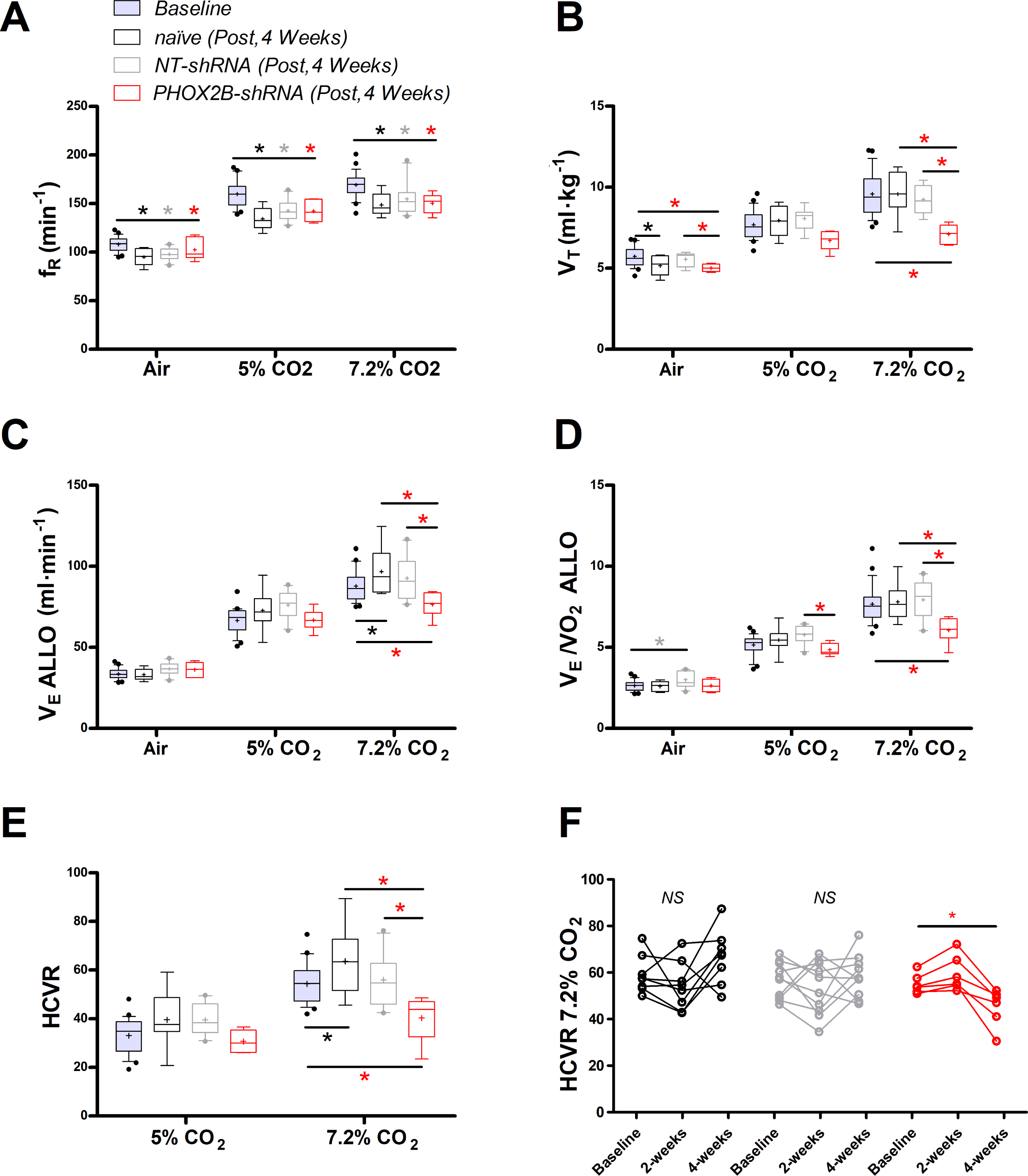
**Respiratory data following 4 weeks post viral shRNA injection**. **(A)** breathing frequency (ƒ_R_,), **(B)** tidal volume (V_T_), **(C)** allometric minute ventilation (V_E_ ALLO), **(D)** convective requirement ratio (V_E_/VO_2_ ALLO), **(E)** hypercapnic ventilatory response (HCVR, absolute change in V_E_ ALLO vs. corresponding room air), **(F)** HCVR at baseline, week 2 and week 4 post-viral injections in naïve (black n=8), non-target control shRNA (NT-shRNA, grey n=10) and PHOX2B-shRNA (PHOX2B- shRNA, red n=6). ƒ_R_ is equally impaired in all experimental group compared to baseline but no treatment effect was observed (A). V_T_ is significantly impaired following RTN injection in PHOX2B- shRNA group compared to baseline pre-surgery (p<0.001), naïve rats p<0.001), and NT-shRNA rats (p=0.002) at 7.2% CO_2_. (B). V_E_ ALLO is impaired in PHOX2B-shRNA rats during exposure to hypercapnia (7.2% CO_2_) compared to baseline (p=0.0025), naïve rats (p=0.007), and NT-shRNA rats (p=0.002). (C). V_E_/VO_2_ ALLO is reduced in PHOX2B-shRNA animals compared to NT-shRNA rats both at 5% (p=0.023) and 7.2% (p=0.004.) CO_2_ (D). HCVR during 7.2% CO_2_ is lower in PHOX2B- shRNA rats compared to baseline (p=0.007), naïve rats (p=0.001), and NT-shRNA rats (p=0.016) (E). Boxplots: median, 1^st^ – 3^rd^ quartiles and 10^th^ – 90^th^ percentiles, outliers = dots, “+” indicates arithmetic mean. Bonferroni post-hoc as indicated. HCVR is significantly impaired only in PHOX2B- shRNA rats 4-weeks post-surgery (One-way ANOVA p=0.007) (F). Black*, different from naïve; Grey*, different from NT-shRNA; Red*, different from PHOX2B- shRNA.

During room air, V_T_ was reduced in naïve rats compared to baseline (baseline: 5.9 ± 0.5 ml·kg^-1^; week 4: 5.2 ± 0.6 ml·kg^-1;^ -12%; p=0.001; Fig. 4B). Moreover, V_T_ in PHOX2B-shRNA rats was reduced compared to NT-shRNA (NT-shRNA: 5.6 ± 0.4 ml·kg^-1^; PHOX2B-shRNA: 5.3 ± 0.4 ml·kg^-1^; -10%; p=0.043) and to the pre-surgery baseline (6.1 ± 0.7 ml·kg^-1^; -18%; p=0.002). No significant changes were observed at 5% CO_2_, although a decrease in V_T_ occurred at 7.2% CO_2_ in PHOX2B-shRNA rats compared to pre-surgery baseline (baseline: 7.7 ± 1.2 ml·kg^-1^; PHOX2B-shRNA: 6.7 ± 0.6 ml·kg^-1^; - 31%; p<0.001), naïve rats (7.9 ± 0.9 ml·kg^-1^; -26%; p<0.001), and NT-shRNA rats (8.1 ± 0.7 ml·kg^-1^; - 23%; p=0.002).

Decreased V_T_ observed for PHOX2B-shRNA led to a ventilation (V_E_ ALLO) impairment during exposure to hypercapnia (7.2% CO_2_, PHOX2B-shRNA vs. NT-shRNA: -17%; p=0.021; PHOX2B-shRNA vs. naïve; -21%; p=0.007; PHOX2B-shRNA vs. baseline, -18% p=0.002; Fig. 4C). Since O_2_- consumption (VO_2_ ALLO) did not change (data not shown) a significant reduction in V_E_/VO_2_ ALLO confirmed alveolar hypoventilation in PHOX2B-shRNA animals compared to NT-shRNA rats both at 5% (NT-shRNA: 5.8 ± 0.5; PHOX2B-shRNA: 4.8 ± 0.4; -16%; p=0.023) and 7.2% CO_2_ (NT-shRNA: 7.9 ± 1.2; PHOX2B-shRNA: 6.1 ± 0.8; -24%; p=0.004; Fig. 4D). Moreover, the HCVR (Fig. 4E) at 7.2% CO_2_ was lower in PHOX2B-shRNA (40.3 ± 9.4 ml·min^-1^) rats compared to the pre-surgery baseline (58.1 ± 8.9 ml·min^-1^; -31%; p=0.015), naïve rats (63.6 ± 14.3 ml·min^-1^; -37%; p=0.008) and NT-shRNA rats (56.0 ± 10.4 ml·min^-1^; -28%; p=0.016). Analysis of individual rats in each treatment group over time showed that the HCVR was only consistently impaired in PHOX2B-shRNA rats between weeks 2 and 4 (Fig. 1F; p=0.007).

Given that previous studies have proposed a role for the RTN in the hypoxic chemoreflex (Barna et al., 2016; Basting et al., 2015; Gourine et al., 2005; Wickström et al., 2004), and we observed impairment of the CO_2_-chemoreflex at 4 weeks post-infection, we investigated whether PHOX2B knockdown would also affect respiratory response to hypoxia (10% O_2_). The 4 week V_E_ ALLO was increased relative to baseline values in naïve (baseline: 62.3 ± 5.2 ml·min^-1^; naïve: 71.0 ± 10.1 ml·min^-1^; +14%), NT-shRNA (baseline: 71.8 ± 11.1 ml·min^-1^; NT-shRNA: 75.8 ± 12.8 ml·min^-1^; +6%) and PHOX2B-shRNA rats (baseline: 65.9 ± 9.8 ml·min^-1^; PHOX2B-shRNA: 79.3 ± 6.4 ml·min^-1^; +29%; p<0.001). However, analysis of the convective requirement ratio (V_E_/VO_2_ ALLO) indicates that changes in hypoxic ventilation were due to metabolic adjustments that occurred in all treatment groups (VE/VO_2_ ALLO naive: 6.5 ± 1.3; NT-shRNA: 6.7 ± 1.7; PHOX2B-shRNA: 6.6 ± 0.6), suggesting that the hypoxic ventilatory response was not affected by PHOX2B knockdown.

### PHOX2B knockdown does not extend to C1 catecholaminergic neurons and does not affect mRNA expression of *Nmb* in RTN neurons

To exclude that the observed chemoreflex impairment was due to unintended knockdown of PHOX2B in adjacent areas of the brain, we quantified the total number of TH^+^/PHOX2B^+^ catecholaminergic C1 neurons in PHOX2B-shRNA rats compared to naïve and NT-shRNA (Fig. 5A,B). No differences were observed in either the TH^+^ cell numbers or in the TH^+^/PHOX2B^+^ neurons between the 3 treatment groups (TH^+^/PHOX2B^+^ cells: naïve: 378.3 ± 23.81; NT-shRNA: 391.7 ± 33.08; PHOX2B-shRNA: 370.2 ± 33.04 cells) (Fig. 5B).

**Figure 5.**
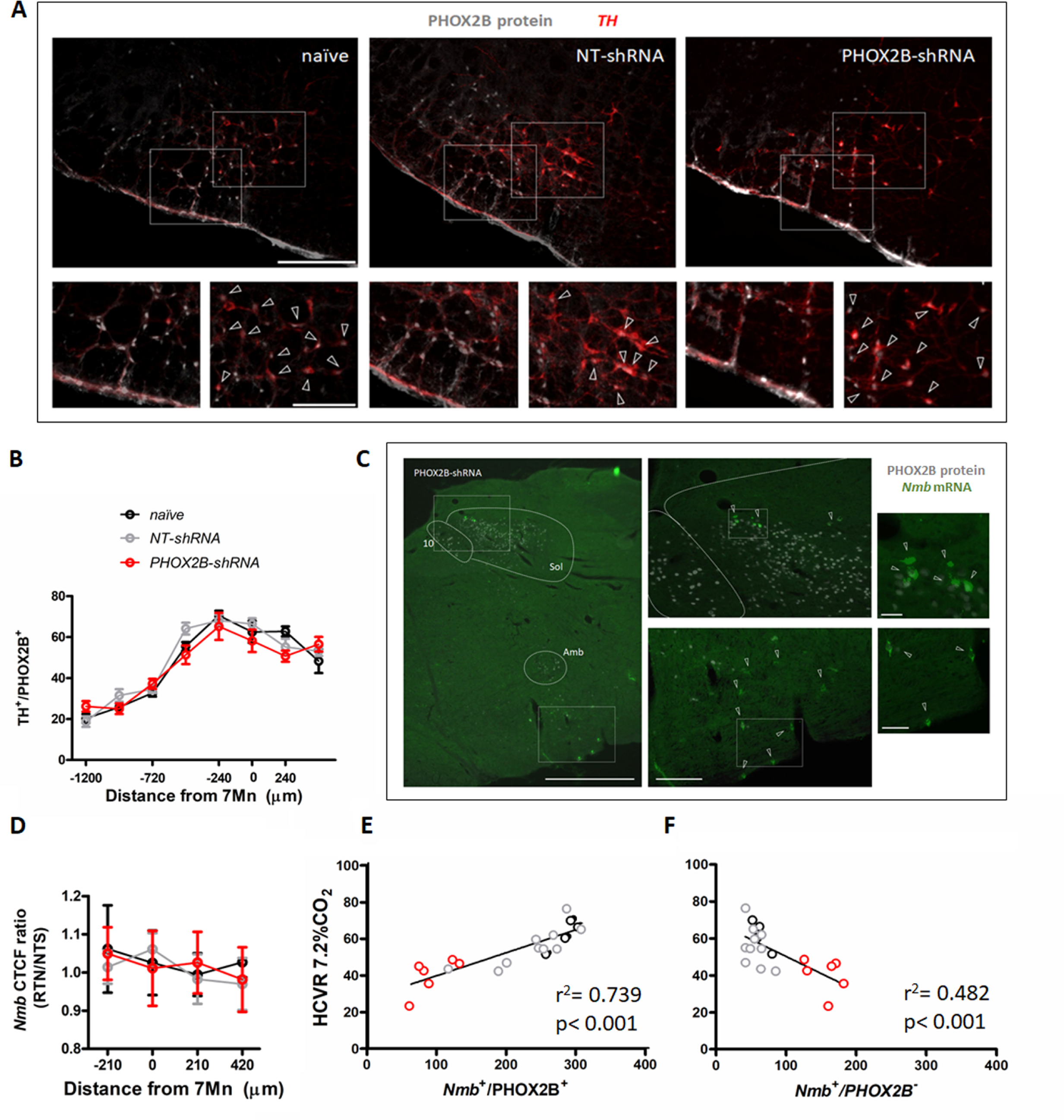
PHOX2B knockdown in RTN neurons does not alter TH or *Nmb* expression but impairs the hypercapnic ventilatory response. (A) PHOX2B protein (white) and TH (red) expression in C1 neurons of naïve (left), NT-shRNA (middle), and PHOX2B-shRNA (right) rats. Magnified view at the bottom. Arrowheads indicate the colocalization of PHOX2B and TH protein in C1 neurones cells. Scale bar = 400μm (top figures), 150μm (bottom figures). **(B)** No differences are observed in the rostral-caudal distribution of TH^+^/PHOX2B^+^ catecholaminergic C1 neurons cells caudal to the RTN between naïve (black, n=4), NT-shRNA (grey, n=10) and PHOX2B-shRNA (red, n=6) rats. **(C)** PHOX2B protein (white) and *Nmb* mRNA (green) expression in NT-shRNA rat at the level of NTS and RTN regions. Magnified view on the left. Arrowheads indicate cells with *Nmb* mRNA expression. Scale bar = 400μm (right figures), 150μm (left figures). **(D)** Quantification of single cells *Nmb* mRNA fluorescence intensity along the rostro-caudal extension of RTN in naïve (black, n=4), NT-shRNA (grey, n=10) and PHOX2B-shRNA (red, n=6) rat calculated as average ratio between RTN and NTS cells shows no difference between treatment groups. Data are shown as average cell fluorescence value at different rostro-caudal levels. **(D-E)** X-Y plot of HCVR during 7.2% CO_2_ exposure relative to the number of *Nmb*^+^/PHOX2B^+^ (D, slope is different from “0” at p<0.001; r^2^=0.739) and *Nmb*^+^/PHOX2B^-^ (E, p<0.001 for difference between slopes; r^2^=0.482) in the RTN.

Neuromedin B is currently considered the most selective anatomical marker for chemosensitive RTN neurons (Shi et al., 2017) and was used in this study to identify RTN location, cell numbers and relative PHOX2B expression. Since changes in the level of the transcription factor PHOX2B could regulate the expression of its target genes, we investigated whether *Nmb* is a target gene of PHOX2B and it is affected by PHOX2B knockdown. We assessed *Nmb* expression in individual RTN cells of PHOX2B-shRNA rats and compared its cellular expression to both naïve and NT-shRNA rats (Fig 5B). Neuromedin B mRNA expression was calculated as total fluorescence intensity (CTCF) ratio in single cells of the RTN relative to the single cell *Nmb* expression measured at the level of NTS *Nmb* cells, which served as an internal control (Fig. 5D). We observed no changes in relative fluorescence of *Nmb* in RTN neurons across the 3 groups (p=0.608), suggesting that *Nmb* levels in RTN neurons are not affected by PHOX2B knockdown.

To better analyse the effect of PHOX2B knockdown on the attenuation of the HCVR, we evaluated the correlation between the number of *Nmb*^+^/PHOX2B^+^ or Nmb^+^/PHOX2B^-^ cells in the RTN and the resulting HCVR using linear regression (Fig. 5E,F). Our results indicate that there is a significant correlation between HCVR and number of Nmb^+^/PHOX2B^+^ (r^2^ _=_ 0.739 p<0.001; Fig. 5E), and Nmb^+^/PHOX2B^-^ (r^2^ _=_ 0.482 p<0.001 Fig. 5F) cells, suggesting that the number of PHOX2B expressing cells in the RTN is a good predictor of the chemoreflex response. Moreover, the nature of the correlation between PHOX2B^-^ cells and HCVR is the opposite of PHOX2B^+^, another important indicator that PHOX2B protein may contribute to the CO_2_ sensing and consequently, the reduction of PHOX2B protein impairs the CO_2_-chemoreflex.

### Gpr4 and Task2 mRNAs expression is affected by PHOX2B Knockdown

We next examined the mRNA expression of *Gpr4* and *Task2*, two important pH sensors for the central respiratory chemoreflex response in RTN neurons (Gestreau et al., 2010; Guyenet et al., 2016; Kumar et al., 2015). As previously reported, the two pH sensors are expressed in the *Nmb* cells of the RTN in partially overlapping populations of chemosensory neurons (Gestreau et al., 2010; Kumar et al., 2015). In naïve rats 91.6 ± 11.2 % of *Nmb* cells were *Gpr4* positive, and 76.6 ± 14.8 % of *Nmb* cells were also *Task2* positive (n=4 rats). These values were not significantly different in either NT-shRNA or PHOX2B shRNA rats (NT-shRNA: 90.0 ± 15.7% *Nmb^+^/Gpr4^+^* and 76.2 ± 21.8% *Nmb^+^/Gpr4*^+^, n=10 rats; PHOX2B-shRNA: 89.4 ± 18.3% *Nmb^+^/Gpr4^+^* and 75.6 ± 27.8% *Nmb^+^/Gpr4^+^*, n=6 rats), suggesting that the fraction of *Nmb* cells expressing the two pH sensors were not affected by the PHOX2B shRNA treatment.

Because it is not yet known whether PHOX2B controls the expression of GPR4 AND TASK2, we quantified their single cell mRNA expression (CTCF) in *Nmb* cells of the RTN. Averaged cell fluorescence for *Gpr4* and *Task2* mRNA in PHOX2B-shRNA rats was significantly reduced compared to naïve and NT-shRNA rats (*Gpr4*: -34.4 ± 4.79% vs naïve, -27.9 ± 5.71% vs NT-shRNA p<0.001; *Task2*: -39.0 ± 3.02% vs naïve, -34 ± 2.62% vs NT-shRNA p<0.001; Fig. 6B,D). To better understand whether this reduction was ascribed to the specific loss of PHOX2B expression, we compared their single cell mRNA expression levels between *Nmb*^+^/PHOX2B^-^ and *Nmb*^+^/PHOX2B^+^ neurons in PHOX2B-shRNA rats. Both *Gpr4* and *Task2* mRNA expression was reduced in *Nmb*^+^/PHOX2B^-^ cells compared to *Nmb*^+^/PHOX2B^+^ (*Gpr4*: -63.8 ± 3.9%; p= 0.022; *Task2*: -63.2 ± 2.77%; p= 0.029; Fig. 6C,E). Our data suggest that PHOX2B knockdown not only causes a reduction in the HCVR but also a reduction in *Gpr4* and *Task2* mRNA levels in cells that are affected by PHOX2B knockdown.

**Figure 6.**
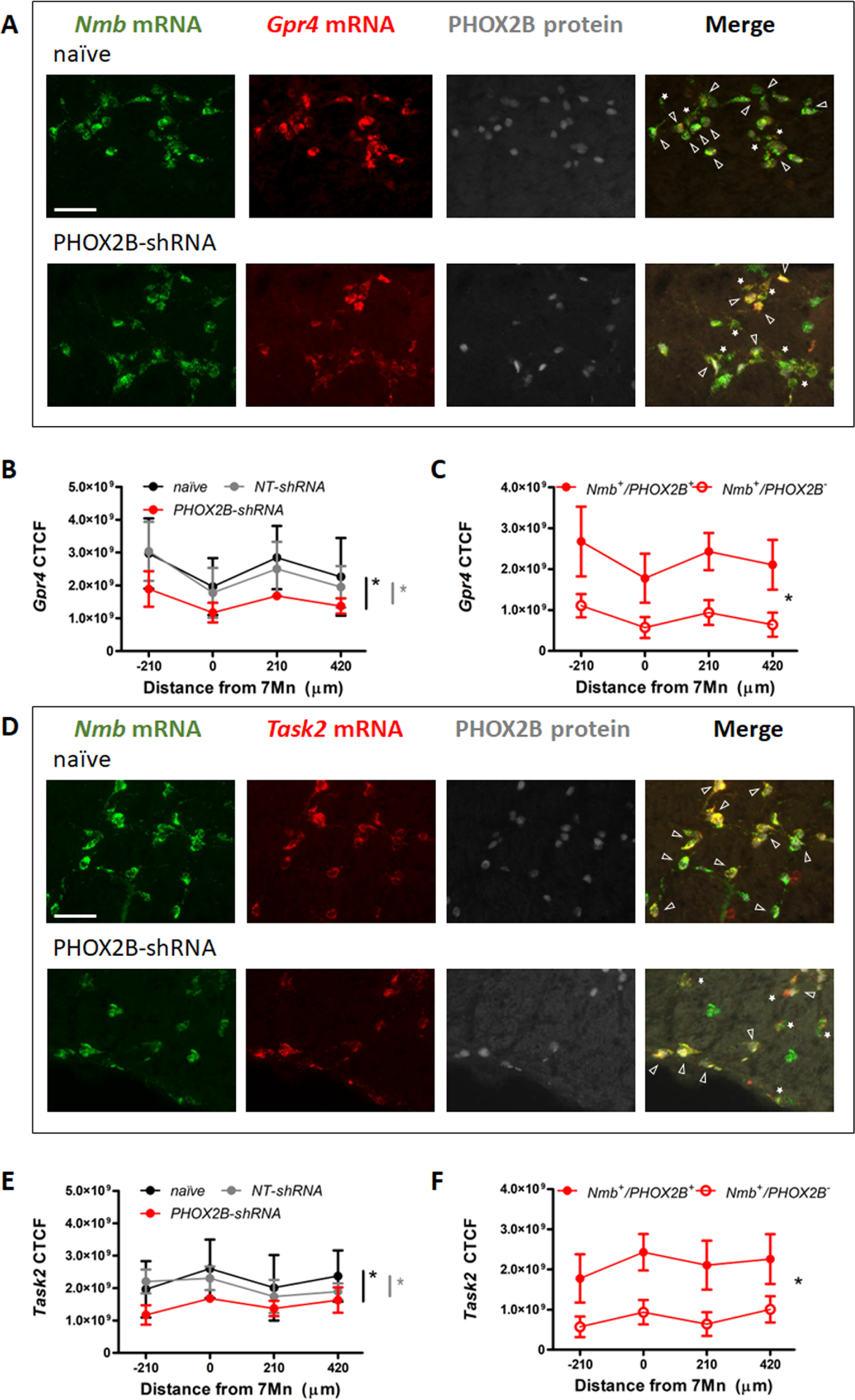
*Gpr4* and *Task2* mRNA expression within the RTN area in naïve, and PHOX2B-shRNA injected rats 4 weeks post viral shRNA injection. (A,D) *Nmb* (green), *Gpr4*, *Task2* mRNA (red) and PHOX2B protein (grey) expression in RTN neurons in naïve (top) and PHOX2B-shRNA (bottom) rats. Scale bar = 150 μm. Arrowheads indicate colocalization of *Gpr4* (A) and *Task2* (D) with *Nmb*^+^/PHOX2B^+^ neurons. Asterisks indicate colocalization of *Gpr4* (A) and *Task2* (D) with *Nmb*^+^/PHOX2B^-^ neurons. **(B,E)** Quantification of single cells *Gpr4* and *Task2* mRNA fluorescence intensity along the rostro-caudal extension of RTN in naïve (black), NT-shRNA (grey) and PHOX2B- shRNA (red) rats. Black*, different from naïve, Grey*, different from NT-shRNA (**C and F**) Quantification of single cells *Gpr4* and *Task2* mRNA fluorescence staining intensity along the rostro-caudal extension of RTN in Nmb^+^/PHOX2B^+^ (red filled dot) and Nmb^+^/PHOX2B^-^ (red empty dot) neurons in PHOX2B-shRNA rats. Data are shown as average cell fluorescence value at different rostro-caudal levels of the RTN. Mean corrected total cell fluorescence (CTCF) value ± SEM combined (naïve, n=4; NT-shRNA, n=10; PHOX2B-shRNA, n=6) (see methods for detail). One- way ANOVA repeated measures.

## DISCUSSION

The role of the transcription factor PHOX2B in the adult brain is still not fully understood. In this study we reduced the expression of PHOX2B in the key chemosensory structure of the RTN with a selective viral shRNA approach to test whether a knockdown of PHOX2B expression in these neurons negatively impacts ventilation. Two weeks after viral infection we observed a modest reduction of PHOX2B/*Nmb* expressing neurons in the RTN but no significant changes in basal V_E_ or in the CO_2_ chemoreflex. Four weeks after shRNA infection we observed a further reduction in the expression of PHOX2B in *Nmb* neurons and a significant reduction of the CO_2_ chemoreflex, while ventilation in normoxia or hypoxia was unaffected. Furthermore, the level of expression of *Gpr4* and *Task2* mRNAs, the two proton sensors responsible for the RTN neurons chemosensory properties were significantly reduced in RTN neurons that lacked PHOX2B protein expression, whereas *Nmb* mRNA expression level was unaffected by PHOX2B knockdown. These results suggest that PHOX2B reduction affects the transcriptional activity of *Nmb* neurons in the RTN and decreases the CO_2_ response, possibly by reducing the expression of key proton sensors in the RTN.

### The role of PHOX2B in development and in the adult brain

PHOX2B is a transcription factor with an important role in the development and differentiation of neuronal structures in both the central and peripheral nervous system. Genetic loss of *Phox2b* expression in mice shows that PHOX2B is essential for the correct development of all autonomic ganglia in sympathetic, parasympathetic, and enteric nervous system and the distal ganglia of the facial (VII), glossopharyngeal (IX), and vagus (X) cranial nerves (Pattyn et al., 1999), in addition to carotid bodies and the solitary tract nucleus (Dauger, 2003). Furthermore, PHOX2B is involved in the formation of all branchial and visceral hindbrain motor neurons and its absence during the embryonic period impairs the development of motoneurons of the facial, trigeminal, ambiguus and dorsal motor nucleus of the vagal nerve. PHOX2B is also necessary for the specification and differentiation of the noradrenergic phenotype in multiple centres in the brain (Brunet & Pattyn, 2002; Pattyn et al., 1999, 2000a).

Interestingly, the expression of PHOX2B persists into adulthood in selected neuronal populations, such as the ones involved in peripheral and central chemoreception, i.e., the carotid bodies, the solitary tract nucleus and the RTN, in addition to neurons in the dorsal motor nucleus of the vagus, the area postrema, the nucleus ambiguus, C1 catecholaminergic neurons and other scattered neurons throughout the brainstem (Stornetta et al., 2006). Such widespread expression within the adult brain suggests an important transcriptional role in neuronal survival and/or function. With the exception of its role in maintaining the noradrenergic phenotype, its function in the postnatal brain is unknown, in part because a full investigation of its transcriptional activity in neurons has not yet been performed.

Heterozygote polyalanine expansion mutations in the exon 3 of *PHOX2B* are responsible for the majority of cases of CCHS, a rare genetic disease that affect the autonomic nervous system and central CO_2_ chemoreflexes (Amiel et al., 2003; Di Lascio et al., 2021; Weese-Mayer et al., 2017). While studies in transgenic mice expressing the mutated PHOX2B protein suggest that the RTN may not form or be severely impaired (Dubreuil et al., 2008, 2009; Ramanantsoa et al., 2011), the function of the wild-type protein in these neurons has never been tested beyond the post-natal period. Thus, in order to determine whether PHOX2B is necessary for the survival and/or chemosensory function of adult PHOX2B^+^/*Nmb*^+^ RTN neurons *in vivo*, we used a shRNA viral approach to progressively knockdown the PHOX2B protein in these neurons.

### Modest reduction of PHOX2B in RTN does not impair ventilation

Although the PHOX2B shRNA approach was not selective for *Nmb* neurons, we used small volumes and localized injections to target the RTN. We identified RTN neurons by the mRNA expression of the neuropeptide NMB (Shi et al., 2017) and observed a reduced number in both total *Nmb* and *Nmb*^+^/PHOX2B^+^ neurons in PHOX2B shRNA treated rats compared to naïve and NT-shRNA rats two weeks after viral infection. Interestingly, despite the small reduction in total *Nmb* cells of the RTN in both NT-shRNA and PHOX2B-shRNA injected rats, possibly due to some off-target effects due to activation of innate immunity or saturation of the microRNA pathway (Mcbride et al., 2008; Van Gestel et al., 2014), no significant impairment in ventilation was observed. Even though PHOX2B may be critical for chemosensory function in RTN neurons, the lack of significant ventilatory impairment with a modest reduction of PHOX2B expressing neurons is not surprising. In our previous study (and the ones of others), in which RTN chemosensory neurons were lesioned with Substance P saporin toxin, CO_2_-chemoreflex impairment became significant (∼30% V_E_ Allo reduction vs. pre-surgical baseline) only when *Nmb* neurons of the RTN (*Nmb*^+^/PHOX2B^+^ and *Nmb*^+^/PHOX2B^-^ neurons) were reduced by ∼63% compared to naïve controls (medium lesion range: 50%-81% loss of *Nmb* neurons; Janes et al., 2024; Souza et al., 2023), therefore, even though loss of PHOX2B severely altered the chemosensitive function in a fraction of *Nmb* cells, we would not expect significant changes in ventilation only with a <50% RTN cell impairment or loss.

### Four weeks of PHOX2B knockdown impairs central CO_2_ chemoreception

Four weeks post- shRNA viral injection, the fraction of *Nmb*^+^ cells expressing PHOX2B was further reduced by 67% compared to naïve and by 61% compared to NT-shRNA rats. PHOX2B knockdown was also restricted to RTN neurons, as adjacent C1 TH^+^ neurons did not show any change in number of TH^+^/PHOX2B^+^ expressing cells, although we can’t exclude that some C1 cells may have been infected, and their relative PHOX2B expression levels were slightly reduced. To support the lack of significant alterations associated with the possible loss of C1 function was the lack of significant changes in the hypoxic response that has been shown to be dependent on C1 neurons (Malheiros-Lima et al., 2017).

Although we did not specifically quantify the relative mRNA expression change in intracellular *Phox2b* levels within infected neurons, we measured the success of PHOX2B knockdown by assessing the overall change in the number of *Nmb* cells that had no detectable levels of PHOX2B protein in the nuclei. Thus, it is possible that this method led us to underestimate the success of the overall PHOX2B shRNA knockdown in the RTN, as some RTN neurons may still have detectable levels of PHOX2B, albeit decreased.

In PHOX2B-shRNA rats we did not observe any respiratory function changes in room air or in hypoxia, similar to RTN lesioning studies (Janes et al., 2024; Souza et al., 2023; Guyenet et al., 2019). When tested in hypercapnia though, PHOX2B-shRNA rats displayed a reduced HCVR compared to both baseline (i.e., pre-surgery), naïve and NT-shRNA rats. The impairment of V_E_ Allo was primarily a result of blunted V_T_, and analysis of the convective exchange ratio (i.e. V_E_/VO_2_ ALLO) confirmed hypoventilation in 5% and 7.2% CO_2_ only for PHOX2B-shRNA rats, suggesting a specific impairment of chemosensitive properties in RTN neurons with PHOX2B knockdown. These results are in line with previous studies in which medium RTN lesions (i.e. loss of >60 % *Nmb*^+^ cells) elicited by saporin toxin injection resulted in a significant CO_2_-chemoreflex impairment (Janes et al., 2024; Souza et al.,2018). Hence, killing RTN neurons or knocking down PHOX2B in *Nmb* cells of the RTN gave comparable results.

Changes in the CO_2_ response following 4 weeks of PHOX2B-shRNA treatment could occur because the reduced expression of the transcription factor PHOX2B causes changes in the transcriptional machinery of RTN neurons and alters transcription of CO_2_/pH sensing proteins or other key elements for their chemosensitive function. These transcriptional changes could also affect neuronal survival. In fact, a small but significant loss of NMB expressing RTN neurons may contribute to the reduction in the CO_2_ response, although multiple studies have demonstrated that significant cell loss must occur in the RTN in order to show a respiratory function impairment (Janes et al., 2024; Souza et al.,2018; Nattie & Li, 2002; Takakura et al., 2008, 2014).

In our experiments, we observed a 35% reduction in total *Nmb*^+^ cells compared to naïve rats and 23% compared to NT-shRNA, possibly due to cell death within the first 14 days post-infection. Interestingly the loss of *Nmb* cells did not increase with time (although the fraction of Nmb^+^/PHOX2B^+^ neurons decreased) and the central chemoreflex at 2 weeks post-surgery was not impaired, suggesting that: i) the observed reduction in overall *Nmb* cell number in RTN occurring in the first 2 weeks is not responsible for the CO_2_ chemoreflex impairment that emerges 4 weeks post-infection; ii) PHOX2B expression is most likely not necessary for neuronal survival of adult *Nmb* neurons of the RTN; iii) a large reduction in *Nmb* neurons expressing PHOX2B^+^ is necessary to impair the HCVR (as we observed at 4 weeks). The interpretation that PHOX2B expression contributes positively to the chemoreflex is further supported by our linear regression analysis showing that *Nmb*^+^/PHOX2B^+^ cell number was the best predictor of the HCVR (r^2^ = 0.739). The initial cell loss, which we observed also in NT-shRNA, and more prominently with larger injection volume of virus, may be attributed to some inherent cell death associated with the surgical procedure (although not observed in similar studies performed by the same investigators, Janes et al., 2024) or, more likely, with the potential off-target toxic effects associated with shRNA procedures (Mcbride et al., 2008; Van Gestel et al., 2014).

### PHOX2B knockdown reduces mRNA expression of proton sensors in the Nmb cells of the RTN

Current theories on central respiratory chemosensitivity postulate that changes in CO_2_/pH in the RTN are detected through two pH sensors, the proton-activated receptor GPR4 and the pH- sensitive K^+^ channel TASK2 that are expressed in partially overlapping populations of NMB expressing RTN neurons (Kumar et al., 2015). Because it is not known whether PHOX2B has any influence on the expression of TASK2, GPR4 or even the expression of the neuropeptide NMB used to anatomically identify RTN neurons, we quantified their mRNA at cellular levels (calculated as CFTC) and observed no changes in relative fluorescence of *Nmb* mRNA in the RTN relative to *Nmb* levels in unrelated cells in the NTS, suggesting that *Nmb* expression is not affected by PHOX2B shRNA knockdown and that *Nmb* is most likely not a target gene of PHOX2B in adult RTN neurons. Furthermore, we determined the mRNA expression levels of *Task2* and *Gpr4* in both *Nmb*^+^/PHOX2B^+^ and Nmb^+^/PHOX2B^-^ neurons in shRNA rats. Our results indicate that although the fraction of *Nmb* cells of the RTN expressing *Gpr4* and *Task2* did not change across treatment, the levels of *Gpr4* and *Task2* in *Nmb* neurons of PHOX2B-shRNA rats was reduced compared to the naïve and NT-shRNA groups. Furthermore, there was a significant reduction of both *Gpr4* and *Task2* levels within *Nmb*^+^/PHOX2B^-^ cells compared to *Nmb*^+^/PHOX2B^+^ cells. Because no good antibodies are currently available to detect protein levels of either NMB, GPR4 or TASK2, our results are only based on changes in mRNA levels, thus we can only speculate that a reduction in *Gpr4* and *Task2* mRNA would translate in a reduction in the protein levels and consequent reduction of RTN neurons chemosensitive properties. Direct electrophysiological recordings in RTN neurons will be able to address the changes in CO_2_/pH sensitivity of RTN neurons at cellular level.

### PHOX2B knockdown may impair CO_2_-sensing through additional transcriptional targets

Since loss of PHOX2B reduces expression of TH and dopamine beta hydroxylase enzymes in catecholaminergic neurons *in vivo* and *in vitro* (Fan et al., 2011), it is possible that in NMB cells of the RTN, reduction of PHOX2B alters the transcriptional machinery of other proteins (in addition to GPR4 and TASK2) and impairs their function in central chemoreception. For example, a change in the expression of enzymes, neurotransmitters, receptors, and ion channels in RTN neurons could affect the excitatory signal transmission to preBötzinger Complex and the respiratory network specifically during CO_2_ challenges (Guyenet et al., 2016). Further studies will be needed to address and identify additional transcriptional changes induced by PHOX2B knockdown *in vivo*.

An interesting observation from our histological data was the reduction in the overall number of *Nmb* cells at two- and four-weeks following PHOX2B shRNA viral injections. Although some cell death may be associated with either surgical procedures or off-target effects associated with the shRNA approach (Mcbride et al., 2008; Van Gestel et al., 2014), it is intriguing to speculate that loss of PHOX2B expression may have some effects on viability of RTN neurons *in vivo*. Even though this is a possibility, especially with an even more severe knockdown, we did observe a large proportion of *Nmb* cells devoid of PHOX2B protein expression, suggesting that absence of PHOX2B is compatible with neuronal survival, at least in our experimental time frame.

### Relevance to CCHS and its pathogenic mechanisms

Our results contribute to further our understanding on potential pathogenic mechanisms in CCHS (Di Lascio et al., 2018). Homozygous knockout mice for PHOX2B die during gestation (Pattyn et al., 1999, 2000) demonstrating a key role of this transcription factor in cellular proliferation, migration and differentiation during embryonic development, whereas heterozygotes for PHOX2B survive birth and live into adulthood, but display apneas (Durand et al., 2005) and impairment of the hypoxic and hypercapnic ventilatory responses in the neonatal period (Dauger, 2003). Here we show that, independent of the PHOX2B role during development, PHOX2B is still required to maintain proper CO_2_ chemoreflex responses in the adult brain.

Heterologous expression of the mutant PHOX2B alters development of RTN in mice and impairs their chemoreflex response (Dubreuil et al., 2008, 2009). The effects of the expression of the mutant PHOX2B on wild-type PHOX2B protein and its transcriptional activity is not known *in vivo* yet, although *in vitro* data indicates that the mutated PHOX2B protein alters wild-type PHOX2B conformation and its ability to form homo- and hetero-dimers (Di Lascio et al., 2016; Trochet et al., 2005, 2008), its affinity for DNA and coactivators, as well as its degradation rate (Nagashimada et al., 2012; Wu et al., 2009). It has also been shown that mutant PHOX2B causes the formation of cytoplasmic aggregates (Bachetti et al., 2005; Trochet et al., 2005, 2008) and fibrils *in vitro* (Pirone et al., 2019), and dysregulates the transcriptional function of wild-type PHOX2B (Di Lascio et al., 2013; Parodi et al., 2012), thus changing expression of important target genes such as DBH, PHOX2A and TLX2 (Bachetti et al., 2005; Di Lascio et al., 2013; Trochet et al., 2005). Based on this evidence, different CCHS pathogenic mechanisms have been proposed (e.g., PHOX2B loss-of-function, dominant-negative or toxic function of the mutant protein) including gene and cell specific transcriptional dysregulation (Di Lascio et al., 2018, 2021). Interestingly, only a handful of PHOX2B target genes have been identified and the function of PHOX2B and its mutated forms in respiratory control and CCHS pathogenesis is still under investigation.

Our data suggest that, in addition to potential effect of mutated PHOX2B on development and cellular function in CCHS pathogenesis, the expression of wild type PHOX2B has an important role in respiratory control that extends the developmental period and its reduction in CCHS may contribute to the respiratory impairment in this disorder.

## MATERIALS AND METHODS

Experiments were performed using male Sprague–Dawley rats (starting weight range 250-350g) born in the University of Alberta animal facility to pregnant dams obtained from Charles River (Senneville, QC). Rats were housed at the University of Alberta Health Sciences Animal Housing Facility and maintained on a 12-hr dark/light cycle with food and water available *ad libitum*. Handling and experimental procedures were approved by the Health Science Animal Policy and Welfare Committee at the University of Alberta (AUP#461) and performed in accordance with guidelines established by the Canadian Council on Animal Care.

### ShRNA viral injection in the retrotrapezoid nucleus

In order to knockdown Phox2b expression at the level of RTN neurons, we used a mix of two shRNA clones targeting two different sequences of the Phox2b mRNA carried by a non-replicating lentivirus vector (1x10^9^ VP/ml; TRCN000041283: GCCTTAGTGAAGAGCAGTATG and TRCN0000096437: CCTCTGCCTACGAGTCCTGTA; Sigma Aldrich, St. Louis, MA, USA). The shRNAs were designed against the Phox2b mouse sequence (Ref Seq NM_008888) which shares 100% homology with the Phox2b rat sequence (Ref Seq XM_008770167).

Metacam analgesic was administered 1 hour prior to surgery (2mg/kg) to reduce post-surgical pain. Rats were anesthetized with a ketamine/xylazine mixture (100mg/kg + 10mg/kg, respectively; i.p.); the anaesthetic level was assessed by lack of paw pinch reflex and maintenance of a regular breathing rate. Additional anaesthesia was administered as needed. The rat was positioned on a stereotactic apparatus (Kopf Instruments, Tujunga, CA, USA) and a total of four microinjections (200 or 100 nL per injection; two rostro-caudally aligned injections per side) were made lateral to the midline and ventral to the facial motor nucleus under aseptic conditions. Coordinates were as follows in mm from obex (medio-lateral/rostro-caudal/dorso-ventral): ±1.8, +2.0,-3.5; ±1.8,+2.4,-3.6. Injections were made using a glass microelectrode (30 μm diameter tip, Drummond Scientific, PA, USA). Twenty-five rats (8 rats 200nl/injection; 17 rats 100nl/injection) underwent shRNA viral injection (PHOX2B-shRNA) and 23 rats (6 rats 200nl/injection; 17 rats 100nl/injection) received a non-target shRNA viral injection (NT-shRNA; SHC016: MISSION^®^ pLKO.1-puro non-Target shRNA Control Plasmid DNA; Sigma Aldrich, St. Louis, MA, USA) to determine any effects of surgery and viral constructs on respiratory behaviour. Eight naïve rats, receiving no surgery and maintained in the same housing conditions, were used as additional controls for both respiratory function and anatomical experiments. Following each injection, the glass electrodes were left in place for 3-5 minutes to minimize backflow of virus up the electrode track. At the end of the surgery, the incision was sutured, and rats received local anaesthetic bupivacaine (0.1 mL, s.c.) and were treated with metacam analgesic for 72 hours. Rats recovered for 7 days before the next respiratory function testing.

### Data acquisition and analysis of respiratory measurements

Rats were habituated to whole-body plethysmography chambers (Buxco, 5L) once, 3-4 days before baseline recordings. On the day of the experiment, rats were placed in the chamber and testing commenced once the animals were quietly awake (typically 20-30 mins). Gas mixtures were delivered at 1.5 L/min using a GSM-3 (CWE Inc., Ardmore, PA, USA) and monitored using a Gas Analyzer (AD Instruments, Sydney, Australia). Ventilatory parameters were measured during exposure to different air compositions for 8-15 min each. Room air: 21%O_2_ + 0%CO_2_ (balanced with N_2_); Normoxic Hypercapnia: 21%O_2_+ 5%CO_2_; 21%O_2_ + 7.2%CO_2_; Normocapnic Hypoxia: 10.6%O_2_+ 0% CO_2_.

As previously described (Cardani et al., 2022; Janes et al., 2024) respiratory data were analysed from rats during quiet wakefulness in the last 5 min period of each gas composition using the barometric method (open-flow system; Cardani et al., 2022; Janes et al., 2024; Mortola & Frappell, 1998; Seifert et al., 2000). Raw pressure signals were acquired with a Validyne differential pressure transducer connected to a CD15 carrier demodulator (Validyne Engineering, Northridge, CA, USA) and digitized using a Powerlab 8/35 (AD Instruments, Sydney, Australia). Analysis was done using Labchart 8 (v8.1.19, AD Instruments, Colorado Springs, CO, USA) to determine tidal volume (V_T_) and breathing frequency (ƒ_R_). The amplitude of the pressure signal was converted to V_T_ (ml·kg^-1^) using the equations of Drorbaug and Fenn 1955 (Drorbaugh & Fenn, 1955) and calibrated against 1 mL of dry air injected into the empty chamber using a rodent ventilator (Harvard Rodent Ventilator Model 683, Holliston, MA, USA ƒ = 50, 75, 100, 150 beats·min^-1^). Humidity and chamber temperature were monitored using a RH-300 water vapour analyzer (Sable Systems, Las Vegas, NV, USA) and a HPR Plus Handheld PIT Tag reader (Biomark, Boise, ID, USA) respectively, depending on the setup. Rectal temperature was read through the temperature probes of a Homeothermic Monitor (507222F, Harvard Apparatus, Holliston, MA, USA). Minute ventilation (^V̇^_E_) was calculated as ƒ_R_ x V_T_ and expressed as ml·min^-1^·kg^-1^. Rats gained, on average, 160 ± 4g over the experimental paradigm (naive: 158 ± 19g; NT-shRNA: 202 ± 25g; PHOX2B-shRNA: 133 ± 7g); we therefore applied an allometric correction to ^V̇^_E_ to account for non-linear effects of weight gain (V_E_ ALLO) (Mortola et al., 1994).

Metabolic parameters VO_2_ and V_E_/VO_2_ were calculated by pull-mode indirect calorimetry and allometrically scaled to be consistent with ventilation (Mortola et al., 1994). The % composition of dry gas flowing into, and out of the recording chamber was measured by using an AD Instruments Gas Analyzer (Colorado Springs, CO) and applying the equations of Depocas and Hart (Depocas F, Hart JS, 1957) and Lighton (Lighton, 2021).

Weight measurements were taken prior to every recording session. Rats breathing function was measured the week prior to surgery (baseline recording) and weekly following shRNA injections to determine the time course of ventilatory impairment.

Respiratory data were compared for naive, NT-shRNA and PHOX2B-shRNA pre- and post-injection using a mixed factor ANOVA (independent factor = treatment, repeated measures factor = pre- and post-lesion; Fig.1,4). Post-hoc analysis used the Bonferroni correction factor, with p<0.05 considered to be significant. Ventilatory data are reported as mean ± standard deviation (SD).

### In Situ hybridization (RNAScope) and immunofluorescence

Either 2 or 4-weeks after viral injection, rats were transcardially perfused with saline 0.9% NaCl followed by 4% paraformaldehyde (PFA) and the brains were post-fixed in 4% PFA overnight and cryoprotected in 30% sucrose in 1X Phosphate Buffered Saline (PBS). Brains were then frozen in O.C.T compound (Fisher Scientific) and sectioned on a cryostat (MODEL CM1950, Leica Biosystems, Buffalo grove, IL, USA) at 30 µm and stored in cryoprotectant buffer at -20°C until processing. Sections were mounted on slides for combined RNAScope® in situ hybridization (Advanced Cell Diagnostics-ACD Bio, Newark CA, USA) and immunofluorescence assay and processed as previously detailed (Biancardi et al., 2021; Cardani et al., 2022). The mRNA expression for *Neuromedin B* (*Nmb)* was used as marker of RTN CO_2_-sensing neurons (Shi et al., 2017). Slides were incubated with probes for *Nmb* (Rn-NMB-C2 #494791-C2, ACDBio, Newark, CA, USA), *Phox2b* (Rn-Phox2b-O1-C1 #1064121-C1, ACDBio), G-protein-coupled receptor 4 (*Gpr4*) (Mm-Gpr4-C1, #427941, ACDBio), and potassium channel, subfamily K, member 5 (*Kcnk5* or *Task- 2*) (Mm-Kcnk5-C3, #427951-C3, ACDBio) for 2 hr at 40°C. Gpr4 and Kcnk5 probes are design against the mouse mRNA sequence (Ref Seq NM_175668.4 and NM_021542.4, respectively). However, the alignment of the mouse mRNA sequences with those of rat (GPR4: Ref Seq NM_001025680.1; TASK2: Ref Seq NM_1039516.2) showed an identity of 94% and 96%, respectively, at the level of the target region (Base Pairs 1030-150) (BLAST ^®^ program, National Library of Medicine, National Center for Biotechnology Information, NIH). In parallel, two sections/rat were treated with positive (low copy housekeeping gene), and negative (non-specific bacterial gene) control probes provided by ACDBio. Finally, slides were processed using the RNAScope Multiplex Fluorescent Assay kit V2 (ACDBio) according to the manufacturer’s instructions. The probes were visualized using Opal 520 and Opal 570 reagent (1:500 and 1:1000, respectively; PerkinElmer, Woodbridge, ON, CA). PHOX2B immunoreactivity was detected using mouse monoclonal PHOX2B antibody (B-11: sc-376997, 1:100, Santa Cruz Biotechnology) incubated overnight in 0.3% TritonX-100, 1%NDS in PBS followed by donkey CY5-conjugated anti-mouse IgG (1:200; Jackson Immuno Research Laboratories Inc) in PBS + 1% NDS for 2 hours. Slides were then washed in PBS and cover-slipped with Fluorosave mounting media (EMD Millipore).

We also performed staining for tyrosine hydroxylase (TH) to identify and quantify C1 cells (TH^+^/PHOX2B^+^) following shRNA injection. Briefly, perfusion and tissue fixation/freezing were done as described above. Floating sections were stained with rabbit anti TH antibody (AB152,1:1000, Millipore SIGMA, USA), and mouse anti PHOX2B (B-11: sc-376997, 1:100, Santa Cruz Biotechnology). Antibodies were visualized using donkey CY3 (TH) and CY5 (PHOX2B) conjugated IgG (1:200; Jackson Immuno Research Laboratories Inc). The sections were washed in PBS and mounted on slides and cover-slipped with Fluorosave mounting media.

### Cell counting, imaging and data analysis

To quantify *Nmb*+/PHOX2B- and *Nmb*+/PHOX2B+ neurons within the RTN region, we analysed one every seven sections (210 µm interval; 8 sections/rat in total) along the rostrocaudal distribution of the RTN on the ventral surface of the brainstem and compared total bilateral cell counts of PHOX2B-shRNA rats with non-target control (NT-shRNA) and naïve rats. Cells that expressed *Nmb* and *Phox2b* mRNAs but did not show co-localization with PHOX2B protein were considered *Nmb*+/PHOX2B-.

The Corrected Total Cell Fluorescence (CTCF) signal for *Nmb, Gpr4* and *Task2* mRNAs was quantified as previously described (Cardani et al., 2022; McCloy et al., 2014). Briefly, a Leica TCS SP5 (B-120G) Laser Scanning Confocal microscope was used to acquire images of the tissue. Exposure time and acquisition parameters were set for the naïve group and kept unchanged for the entire dataset acquisition. The collected images were then analysed by selecting a single cell at a time and measuring the area, integrated density and mean grey value (McCloy et al., 2014). For each image, three background areas were used to normalize against autofluorescence. We used 4 sections/rat (210 µm interval) to count *Nmb*, *Gpr4* and *Task2* mRNA CTCF in the core of the RTN area where several *Nmb* cells could be identified. For each section, two images were acquired with a 20*×* objective, so that at least fifty cells per tissue sample were obtained for the mRNA quantification analysis. To evaluate changes in *Nmb* mRNA expression levels following PHOX2B knockdown at the level of the RTN, we compared, the fluorescence intensity of each RTN *Nmb* cell (223.2 ± 37.1 cells/animal) with the average fluorescent signal of *Nmb* cells located dorsally in the NTS (4.3 ± 1.2 cells/animal) (*Nmb* CTCF ratio RTN/NTS) as we reasoned that the latter would not be affected by the shRNA infection and knockdown.

To quantify *Gpr4* and *Task2* mRNA expression in *Nmb* cells of the RTN, we first quantified single cell CTCF for either *Gpr4* (200.7 ± 13.2 cells/rat) or *Task2* (169.6 ± 10.3 cells/rat) mRNA in *Nmb* cells of the RTN in the 3 experimental groups (naïve, NT shRNA and PHOX2B shRNA) independent of their PHOX2B expression. We then compared CTCF values of *Gpr4* and *Task2* mRNA between *Nmb*^+^/PHOX2B^+^ and *Nmb*^+^/PHOX2B^-^ RTN neurons in PHOX2B-shRNA rats to address changes in their mRNA expression induced by PHOX2B knockdown.

To assess whether shRNA knockdown affected either the number or the expression of PHOX2B in TH expressing C1 neurons, we counted the number of TH^+^/PHOX2B^+^ and TH^+^/PHOX2B^-^ cells along the ventrolateral medulla (8 sections/rat; 240 µm interval) using an EVOS fluorescent microscope (ThermoFisher Scientific).

Digital colour photomicrographs were acquired using a Leica TCS SP5 (B-120G) Laser Scanning Confocal microscope. Image J (version 1.54f; National Institutes of Health, Bethesda, MD, USA) and Excel (Microsoft Office 365 for Windows) were used for cell counting and the measurements of fluorescence intensity. Statistical analysis of cell counts, and fluorescent signals was made using one- way ANOVA (Fig. 2C, 3C, S1B) and repeated-measures ANOVA (Fig 5B, D and 6) with GraphPad Prism 8 Software (GraphPad Software, Inc., San Diego, CA, USA). Linear regressions to determine the effect of cell counts (*Nmb*^+^/PHOX2B^+^, *Nmb*^+^/PHOX2B^-^, total cells) on chemoreflex magnitude were run using a combined dataset for naive, NT-shRNA and PHOX2B-shRNA rats (SPSS version 13.0; Fig.5E-F). *p* values of < 0.05 were considered significant and data are reported as mean ± standard deviation (SD).

## Acknowledgments/ Author contributions

Concept and experimental design, S.P. and S.C.; Data collection, S.C., W.B. and S.P.; Data analysis, S.C. and T.A.J.; Manuscript preparation, S.C., T.A.J. and S.P. S.C. is supported by the Canadian Lung Association BaO Fellowship. T.A.J. is supported by the Canadian Institutes for Health Research (CIHR) Postdoctoral Fellowships. S.P. is supported by a Women and Children’s Health Research Institute Innovation Grant and CIHR Project scheme grant.

---

**Supplementary Fig. 1. (A)** Allometric V_E_ is equally impaired following RTN injection of non-target control (NT-shRNA n=6) and PHOX2B-shRNA (n=8). Asterisks, difference from baseline. **(B)** The number of Nmb^+^/PHOX2B^+^ cells comprising the RTN are reduced in NT-shRNA (n=5), and shRNA rats (n=5) as compared to naive controls (n=4; Asterisks, difference from baseline). The number of Nmb^+^/PHOX2B^-^ cells was unchanged in both surgical treatments compared to baseline.

**Supplementary table 1.** Summary of statistical tests and results presented in the figures.

## REFERENCES

1. Abbott, S. B. G., Stornetta, R. L., Coates, M. B., & Guyenet, P. G. (2011). Phox2b-Expressing Neurons of the Parafacial Region Regulate Breathing Rate, Inspiration, and Expiration in Conscious Rats. 10.1523/JNEUROSCI.3280-11.2011

2. Amiel, J., Laudier, B., Attié-Bitach, T., Trang, H., De Pontual, L., Gener, B., Trochet, D., Etchevers, H., Ray, P., Simonneau, M., Vekemans, M., Munnich, A., Gaultier, C., & Lyonnet, S. (2003). Polyalanine expansion and frameshift mutations of the paired-like homeobox gene PHOX2B in congenital central hypoventilation syndrome. Nature Genetics, 33(4), 459–461. 10.1038/ng1130

3. Bachetti, T., Matera, I., Borghini, S., Duca, M. Di, Ravazzolo, R., & Ceccherini, I. (2005). Distinct pathogenetic mechanisms for PHOX2B associated polyalanine expansions and frameshift mutations in congenital central hypoventilation syndrome. Human Molecular Genetics, 14(13), 1815–1824. 10.1093/hmg/ddi188

4. Barna, B. F., Takakura, A. C., Mulkey, D. K., & Moreira, T. S. (2016). Purinergic receptor blockade in the retrotrapezoid nucleus attenuates the respiratory chemoreflexes in awake rats. *Acta Physiologica (Oxford*, England), 217(1), 80–93. 10.1111/APHA.12637

5. Basting, T. M., Burke, P. G. R., Kanbar, R., Viar, K. E., Stornetta, D. S., Stornetta, R. L., & Guyenet, P. G. (2015). Hypoxia Silences Retrotrapezoid Nucleus Respiratory Chemoreceptors via Alkalosis. 10.1523/JNEUROSCI.2923-14.2015

6. Berry-Kravis, E. M., Zhou, L., Rand, C. M., & Weese-Mayer, D. E. (2006). Congenital central hypoventilation syndrome PHOX2B mutations and phenotype. American Journal of Respiratory and Critical Care Medicine, 174(10), 1139–1144. 10.1164/rccm.200602-305OC

7. Biancardi, V., Saini, J., Pageni, A., Prashaad M., H., Funk, G. D., & Pagliardini, S. (2021). Mapping of the excitatory, inhibitory, and modulatory afferent projections to the anatomically defined active expiratory oscillator in adult male rats. Journal of Comparative Neurology, 529(4), 853– 884. 10.1002/CNE.24984

8. Bourdeaut, F., Trochet, D., Janoueix-Lerosey, I., Ribeiro, A., Deville, A., Coz, C., Michiels, J. F., Lyonnet, S., Amiel, J., & Delattre, O. (2005). Germline mutations of the paired-like homeobox 2B (PHOX2B) gene in neuroblastoma. In Cancer Letters (Vol. 228, Issues 1–2, pp. 51–58). Elsevier. 10.1016/j.canlet.2005.01.055

9. Brunet, J. F., & Pattyn, A. (2002). Phox2 genes - From patterning to connectivity. In Current Opinion in Genetics and Development (Vol. 12, Issue 4, pp. 435–440). 10.1016/S0959-437X(02)00322-2

10. Cardani, S., Janes, T. A., Saini, J. K., Di Lascio, S., Benfante, R., Fornasari, D., & Pagliardini, S. (2022). Etonogestrel Administration Reduces the Expression of PHOX2B and Its Target Genes in the Solitary Tract Nucleus. International Journal of Molecular Sciences, 23(9). 10.3390/ijms23094816

11. Cutsforth-Gregory, J. K., & Benarroch, E. E. (2017). Nucleus of the solitary tract, medullary reflexes, and clinical implications. Neurology, 88(12), 1187–1196. 10.1212/WNL.0000000000003751

12. Dauger, S. (2003). Phox2b controls the development of peripheral chemoreceptors and afferent visceral pathways. Development, 130(26), 6635–6642. 10.1242/dev.00866

13. Depocas, F., & Sanford Hart, J. (n.d.). Use of the Pdzlling Oxygen Analyzer for Memwement of Oxygen conszlm∼tion of Ahmds iti Open-Circuit Systems ad in a Short-Lag, Closed-Cirwit Appma tml.

14. Di Lascio, S., Bachetti, T., Saba, E., Ceccherini, I., Benfante, R., & Fornasari, D. (2013). Transcriptional dysregulation and impairment of PHOX2B auto-regulatory mechanism induced by polyalanine expansion mutations associated with congenital central hypoventilation syndrome. Neurobiology of Disease, 50(1), 187–200. 10.1016/j.nbd.2012.10.019

15. Di Lascio, S., Belperio, D., Benfante, R., & Fornasari, D. (2016). Alanine expansions associated with congenital central hypoventilation syndrome impair PHOX2B homeodomain-mediated dimerization and nuclear import. Journal of Biological Chemistry, 291(25), 13375–13393. 10.1074/jbc.M115.679027

16. Di Lascio, S., Benfante, R., Cardani, S., & Fornasari, D. (2018). Advances in the molecular biology and pathogenesis of congenital central hypoventilation syndrome—implications for new therapeutic targets. Expert Opinion on Orphan Drugs, 00(00), 1–13. 10.1080/21678707.2018.1540978

17. Di Lascio, S., Benfante, R., Cardani, S., & Fornasari, D. (2021). Research Advances on Therapeutic Approaches to Congenital Central Hypoventilation Syndrome (CCHS). Frontiers in Neuroscience, 14. 10.3389/FNINS.2020.615666

18. Di Lascio, S., Benfante, R., Zanni, E. Di, Cardani, S., Adamo, A., Fornasari, D., Ceccherini, I., & Bachetti, T. (2018). Structural and functional differences in PHOX2B frameshift mutations underlie isolated or syndromic congenital central hypoventilation syndrome. 10.1002/humu.23365

19. Drorbaugh, J. E., & Fenn, W. O. (1955). A barometric method for measuring ventilation in newborn infants. Pediatrics, 16(1), 81–87. 10.1542/peds.16.1.81

20. Dubreuil, V., Ramanantsoa, N., Trochet, D., Vaubourg, V., Amiel, J., Gallego, J., Brunet, J.-F., & Goridis, C. (2008). A human mutation in Phox2b causes lack of CO2 chemosensitivity, fatal central apnea, and specific loss of parafacial neurons. Proceedings of the National Academy of Sciences, 105(3), 1067–1072. 10.1073/pnas.0709115105

21. Dubreuil, V., Thoby-Brisson, M., Rallu, M., Persson, K., Pattyn, A., Birchmeier, C., Brunet, J.-F., Fortin, G., & Goridis, C. (2009). Defective Respiratory Rhythmogenesis and Loss of Central Chemosensitivity in Phox2b Mutants Targeting Retrotrapezoid Nucleus Neurons. Journal of Neuroscience, 29(47), 14836–14846. 10.1523/JNEUROSCI.2623-09.2009

22. Durand, E., Dauger, S., Pattyn, A., Gaultier, C., Goridis, C., & Gallego, J. (2005). Sleep-disordered breathing in newborn mice heterozygous for the transcription factor phox2b. American Journal of Respiratory and Critical Care Medicine, 172(2), 238–243. 10.1164/rccm.200411-1528OC

23. Fan, Y., Huang, J., Duffourc, M., Kao, R. L., Ordway, G. A., Huang, R., & Zhu, M. Y. (2011). Transcription factor Phox2 upregulates expression of norepinephrine transporter and dopamine β-hydroxylase in adult rat brains. Neuroscience, 192, 37–53. 10.1016/j.neuroscience.2011.07.005

24. Gestreau, C., Heitzmann, D., Thomas, J., Dubreuil, V., Bandulik, S., Reichold, M., Bendahhou, S., Pierson, P., Sterner, C., Peyronnet-Roux, J., Benfriha, C., Tegtmeier, I., Ehnes, H., Georgieff, M., Lesage, F., Brunet, J.-F., Goridis, C., Warth, R., & Barhanin, J. (2010). Task2 potassium channels set central respiratory CO2 and O2 sensitivity. Proceedings of the National Academy of Sciences, 107(5), 2325–2330. 10.1073/pnas.0910059107

25. Goridis, C., Dubreuil, V., Thoby-brisson, M., & Fortin, G. (2010). Seminars in Cell & Developmental Biology Phox2b, congenital central hypoventilation syndrome and the control of respiration. 21, 814–822. 10.1016/j.semcdb.2010.07.006

26. Gourine, A. V, Llaudet, E., Dale, N., & Spyer, K. M. (2005). Behavioral/Systems/Cognitive Release of ATP in the Ventral Medulla during Hypoxia in Rats: Role in Hypoxic Ventilatory Response. 10.1523/JNEUROSCI.3763-04.2005

27. Guyenet, P. G., Bayliss, D. A., Stornetta, R. L., Ludwig, M. G., Kumar, N. N., Shi, Y., Burke, P. G. R., Kanbar, R., Basting, T. M., Holloway, B. B., & Wenker, I. C. (2016). Proton detection and breathing regulation by the retrotrapezoid nucleus. Journal of Physiology, 594(6), 1529–1551. 10.1113/JP271480

28. Guyenet, P. G., Stornetta, R. L., Souza, G. M. P. R., Abbott, S. B. G., Shi, Y., & Bayliss, D. A. (2019). The Retrotrapezoid Nucleus: Central Chemoreceptor and Regulator of Breathing Automaticity. Trends in Neurosciences, 42(11), 807–824. 10.1016/J.TINS.2019.09.002

29. Harper, R. M., Kumar, R., Macey, P. M., Harper, R. K., & Ogren, J. A. (2015). Impaired neural structure and function contributing to autonomic symptoms in congenital central hypoventilation syndrome. Frontiers in Neuroscience, 9(OCT), 415. 10.3389/FNINS.2015.00415

30. Janes, T. A., Cardani, S., Saini, J. K., & Pagliardini, S. (2024b). Etonogestrel promotes respiratory recovery in an in vivo rat model of central chemoreflex impairment. Acta Physiologica. 10.1111/apha.14093

31. Kang, B. J., Chang, D. A., Mackay, D. D., West, G. H., Moreira, T. S., Takakura, A. C., Gwilt, J. M., Guyenet, P. G., & Stornetta, R. L. (2007). Central nervous system distribution of the transcription factor Phox2b in the adult rat. Journal of Comparative Neurology, 503(5), 627– 641. 10.1002/cne.21409

32. Kumar, N. N., Velic, A., Soliz, J., Shi, Y., Li, K., Wang, S., Weaver, J. L., Sen, J., Abbott, S. B. G., Lazarenko, R. M., Ludwig, M. G., Perez-Reyes, E., Mohebbi, N., Bettoni, C., Gassmann, M., Suply, T., Seuwen, K., Guyenet, P. G., Wagner, C. A., & Bayliss, D. A. (2015). Regulation of breathing by CO2 requires the proton-activated receptor GPR4 in retrotrapezoid nucleus neurons. Science, 348(6240), 1255–1260. 10.1126/science.aaa0922

33. Lighton, J. R. B. (2021). MEASURING METABOLIC RATES : a manual for scientists. https://global.oup.com/academic/product/measuring-metabolic-rates-9780198869320

34. Madani, A., Pitollat, G., Sizun, E., Cardoit, L., Ringot, M., Bourgeois, T., Ramanantsoa, N., Delclaux, C., Dauger, S., Thoby-Brisson, M., Gallego, J., & Matrot, B. (2021). Obstructive Apneas in a Mouse Model of Congenital Central Hypoventilation Syndrome. American journal of respiratory and critical care medicine, 204(10), 1200–1210. 10.1164/rccm.202104-0887OC.

35. Malheiros-Lima MR, Takakura AC, Moreira TS. Depletion of rostral ventrolateral medullary catecholaminergic neurons impairs the hypoxic ventilatory response in conscious rats. Neuroscience. 2017 May 20;351:1–14. doi: 10.1016/j.neuroscience.2017.03.031.

36. Marina, N., Abdala, A. P., Trapp, S., Li, A., Nattie, E. E., Hewinson, J., Smith, J. C., Paton, J. F. R., & Gourine, A. V. (2010). Essential Role of Phox2b-Expressing Ventrolateral Brainstem Neurons in the Chemosensory Control of Inspiration and Expiration. Journal of Neuroscience, 30(37), 12466–12473. 10.1523/JNEUROSCI.3141-10.2010

37. Matera, I. (2004). PHOX2B mutations and polyalanine expansions correlate with the severity of the respiratory phenotype and associated symptoms in both congenital and late onset Central Hypoventilation syndrome. Journal of Medical Genetics, 41(5), 373–380. 10.1136/jmg.2003.015412

38. Mcbride, J. L., Boudreau, R. L., Harper, S. Q., Staber, P. D., Monteys, A. M., Martins, I. S., Gilmore, B. L., Burstein, H., Peluso, R. W., Polisky, B., Carter, B. J., & Davidson, B. L. (2008). Artificial miRNAs mitigate shRNA-mediated toxicity in the brain: implications for the therapeutic development of RNAi. Proceedings of the National Academy of Sciences of the United States of America, 105(15), 5868–5873. 10.1073/pnas.0801775105

39. McCloy, R. A., Rogers, S., Caldon, C. E., Lorca, T., Castro, A., & Burgess, A. (2014). Partial inhibition of Cdk1 in G2 phase overrides the SAC and decouples mitotic events. Cell Cycle, 13(9), 1400– 1412. 10.4161/CC.28401/SUPPL_FILE/KCCY_A_10928401_SM0001.ZIP

40. Mortola, J. P., & Frappell, P. B. (1998). On the barometric method for measurements of ventilation, and its use in small animals.

41. Mortola, J. P., Matsuoka, T., Saiki, C., & Naso, L. (1994). RESPIRATION PHYSIOLOGY Metabolism and ventilation in hypoxic rats: effect of body mass. Respiration Physiology, 97, 225–234.

42. Nagashimada, M., Ohta, H., Li, C., Nakao, K., Uesaka, T., Brunet, J. F., Amiel, J., Trochet, D., Wakayama, T., & Enomoto, H. (2012). Autonomic neurocristopathy-associated mutations in PHOX2B dysregulate Sox10 expression. Journal of Clinical Investigation, 122(9), 3145–3158. 10.1172/JCI63401

43. Nattie, E. E., & Li, A. (1994). Retrotrapezoid nucleus lesions decrease phrenic activity and CO2 sensitivity in rats. Respiration Physiology, 97(1), 63–77. 10.1016/0034-5687(94)90012-4

44. Nattie, E. E., & Li, A. (2002). Substance P-saporin lesion of neurons with NK1 receptors in one chemoreceptor site in rats decreases ventilation and chemosensitivity. The Journal of Physiology, 544(2), 603–616. 10.1113/JPHYSIOL.2002.020032

45. Nobuta, H., Cilio, M. R., Danhaive, O., Tsai, H.-H., Tupal, S., Chang, S. M., Murnen, A., Kreitzer, F., Bravo, V., Czeisler, C., Gokozan, H. N., Gygli, P., Bush, S., Weese-Mayer, D. E., Conklin, B., Yee, S.-P., Huang, E. J., Gray, P. A., Rowitch, D., & Otero, J. J. (2015). Dysregulation of locus coeruleus development in congenital central hypoventilation syndrome. Acta Neuropathologica, 130(2), 171–183. 10.1007/s00401-015-1441-0

46. Parodi, S., Di Zanni, E., Di Lascio, S., Bocca, P., Prigione, I., Fornasari, D., Pennuto, M., Bachetti, T., Ceccherini, I., Zanni, E. Di, Lascio, D., Bocca, P., Prigione, I., Fornasari, D., Pennuto, M., & Bachetti, T. (2012). The E3 ubiquitin ligase TRIM11 mediates the degradation of congenital central hypoventilation syndrome-associated polyalanine-expanded PHOX2B. Journal of Molecular Medicine, 90(9), 1025–1035. 10.1007/s00109-012-0868-1

47. Pattyn, A., Hirsch, M. R., Goridis, C., & Brunet, J. F. (2000). Control of hindbrain motor neuron differentiation by the homeobox gene Phox2b. *Development (Cambridge*, England), 127(7), 1349–1358. 10.1242/DEV.127.7.1349

48. Pattyn, A., Morin, X., & Cremer, H. (1999). The homeobox gene Phox2b is essential for the development of autonomic neural crest derivatives. *399*(May).

49. Pattyn, A., Morin, X., Cremer, H., Goridis, C., & Brunet, J. F. (1997). Expression and interactions of the two closely related homeobox genes Phox2a and Phox2b during neurogenesis. *Development (Cambridge*, England), 124(20), 4065–4075. http://www.ncbi.nlm.nih.gov/pubmed/9374403

50. Paxinos, G., & Watson, C. The rat brain in stereotaxic coordinates, 2007

51. Pirone, L., Caldinelli, L., Di Lascio, S., Di Girolamo, R., Di Gaetano, S., Fornasari, D., Pollegioni, L., Benfante, R., & Pedone, E. (2019). Molecular insights into the role of the polyalanine region in mediating PHOX2B aggregation. FEBS Journal, 286(13), 2505–2521. 10.1111/FEBS.14841

52. Ramanantsoa, N., Hirsch, M.-R., Thoby-Brisson, M., Dubreuil, V., Bouvier, J., Ruffault, P.-L., Matrot, B., Fortin, G., Brunet, J.-F., Gallego, J., & Goridis, C. (2011). Breathing without CO2 Chemosensitivity in Conditional Phox2b Mutants. Journal of Neuroscience, 31(36), 12880– 12888. 10.1523/JNEUROSCI.1721-11.2011

53. Seifert, E. L., Knowles, J., & Mortola, J. P. (2000). Continuous circadian measurements of ventilation in behaving adult rats. Respiration Physiology, 120, 179–183. www.elsevier.com/locate/resphysiol

54. Shi, Y., Stornetta, R. L., Stornetta, D. S., Suna Onengut-Gumuscu, X., Farber, E. A., Turner, S. D., Guyenet, P. G., & Bayliss, D. A. (2017). Neuromedin B Expression Defines the Mouse Retrotrapezoid Nucleus. 10.1523/JNEUROSCI.2055-17.2017

55. Shimokaze, T., Sasaki, A., Meguro, T., Hasegawa, H., Hiraku, Y., Yoshikawa, T., Kishikawa, Y., & Hayasaka, K. (2015). Genotype-phenotype relationship in Japanese patients with congenital central hypoventilation syndrome. Journal of Human Genetics, 60(9), 473–477. 10.1038/jhg.2015.65

56. Souza, G. M. P. R., Kanbar, R., Stornetta, D. S., Abbott, S. B. G., Stornetta, R. L., & Guyenet, P. G. (2018). Breathing regulation and blood gas homeostasis after near complete lesions of the retrotrapezoid nucleus in adult rats. Journal of Physiology, 596(13), 2521–2545. 10.1113/JP275866

57. Souza, G. M. P. R., Stornetta, D. S., Shi, Y., Lim, E., Berry, F. E., Bayliss, D. A., & Abbott, S. B. G. (2023). Neuromedin B-Expressing Neurons in the Retrotrapezoid Nucleus Regulate Respiratory Homeostasis and Promote Stable Breathing in Adult Mice. Journal of Neuroscience, 43(30), 5501–5520. 10.1523/JNEUROSCI.0386-23.2023

58. Stanke, M., Junghans, D., Geissen, M., Goridis, C., Ernsberger, U., & Rohrer, H. (1999). The Phox2 homeodomain proteins are sufficient to promote the development of sympathetic neurons. Development, 126(18), 4087–4094. 10.1242/dev.00165

59. Stornetta, R. L., Moreira, T. S., Takakura, A. C., Kang, B. J., Chang, D. A., West, G. H., Mulkey, D. K., Bayliss, D. A., & Guyenet, P. G. (2006). Expression of Phox2b by Brainstem Neurons Involved in Chemosensory Integration in the Adult Rat. 26(40), 10305–10314. 10.1523/JNEUROSCI.2917-06.2006

60. Takakura, A. C., Barna, B. F., Cruz, J. C., Colombari, E., & Moreira, T. S. (2014). Phox2b-expressing retrotrapezoid neurons and the integration of central and peripheral chemosensory control of breathing in conscious rats. Experimental Physiology, 99(3). 10.1113/expphysiol.2013.076752

61. Takakura, A. C., Moreira, T. S., Stornetta, R. L., West, G. H., Gwilt, J. M., & Guyenet, P. G. (2008). Selective lesion of retrotrapezoid Phox2b-expressing neurons raises the apnoeic threshold in rats. The Journal of Physiology, 586(Pt 12), 2975. 10.1113/JPHYSIOL.2008.153163

62. Tiveron, M. C., Hirsch, M. R., & Brunet, J. F. (1996). The expression pattern of the transcription factor Phox2 delineates synaptic pathways of the autonomic nervous system. The Journal of Neuroscience : The Official Journal of the Society for Neuroscience, 16(23), 7649–7660. 10.1523/JNEUROSCI.16-23-07649.1996

63. Trang, H., Dehan, M., Beaufils, F., Zaccaria, I., Amiel, J., & Gaultier, C. (2005). The French Congenital Central Hypoventilation Syndrome Registry. Chest, 127(1), 72–79. 10.1378/chest.127.1.72

64. Trochet, D., de Pontual, L., Estêvao, M. H., Mathieu, Y., Munnich, A., Feingold, J., Goridis, C., Lyonnet, S., & Amiel, J. (2008). Homozygous mutation of the PHOX2B gene in congenital central hypoventilation syndrome (Ondine’s Curse). Human Mutation, 29(5), 770–770. 10.1002/humu.20727

65. Trochet, D., Hong, S. J., Lim, J. K., Brunet, J. F., Munnich, A., Kim, K. S., Lyonnet, S., Goridis, C., & Amiel, J. (2005). Molecular consequences of PHOX2B missense, frameshift and alanine expansion mutations leading to autonomic dysfunction. Human Molecular Genetics, 14(23), 3697–3708. 10.1093/hmg/ddi401

66. Van Gestel, M. A., Van Erp, S., Sanders, L. E., Brans, M. A. D., Luijendijk, M. C. M., Merkestein, M., Pasterkamp, R. J., & Adan, R. A. H. (2014). shRNA-induced saturation of the microRNA pathway in the rat brain. Gene Therapy, 21(2), 205–211. 10.1038/GT.2013.76

67. Van Limpt, V., Schramm, A., Lakeman, A., Van Sluis, P., Chan, A., Van Noesel, M., Baas, F., Caron, H., Eggert, A., & Versteeg, R. (2004). The Phox2B homeobox gene is mutated in sporadic neuroblastomas. Oncogene, 23(57), 9280–9288. 10.1038/sj.onc.1208157

68. Weese-Mayer, D. E., Berry-Kravis, E. M., Zhou, L., Maher, B. S., Silvestri, J. M., Curran, M. E., & Marazita, M. L. (2003). Idiopathic congenital central hypoventilation syndrome: Analysis of genes pertinent to early autonomic nervous system embryologic development and identification of mutations in PHOX2b. American Journal of Medical Genetics, *123A*(3), 267–278. 10.1002/ajmg.a.20527

69. Weese-Mayer, D. E., Rand, C. M., Zhou, A., Carroll, M. S., & Hunt, C. E. (2017). Congenital central hypoventilation syndrome: A bedside-to-bench success story for advancing early diagnosis and treatment and improved survival and quality of life. In Pediatric Research (Vol. 81, Issues 1–2). 10.1038/pr.2016.196

70. Wickström, H. R., Berner, J., Holgert, H., Hökfelt, T., & Lagercrantz, H. (2004). Hypoxic response in newborn rat is attenuated by neurokinin-1 receptor blockade. Respiratory Physiology and Neurobiology, 140(1), 19–31. 10.1016/j.resp.2004.01.008

71. Wu, H., Su, Y., Hung, C., Hsieh, W., & Wu, K. (2009). Interaction Between PHOX2B and CREBBP Mediates Synergistic Activation : Mechanistic Implications of PHOX2B Mutants Human Mutation. 10.1002/humu.20929

72. Zoccal, D. B., Furuya, W. I., Bassi, M., A Colombari, D. S., Colombari, E., Hildreth, C., & Putnam, R. W. (2014). The nucleus of the solitary tract and the coordination of respiratory and sympathetic activities. 10.3389/fphys.2014.00238

