## Supplementary figures and images for "Knockdown of PHOX2B in the retrotrapezoid nucleus reduces the central CO_2_ chemoreflex in rats"

### supplemental figure 1

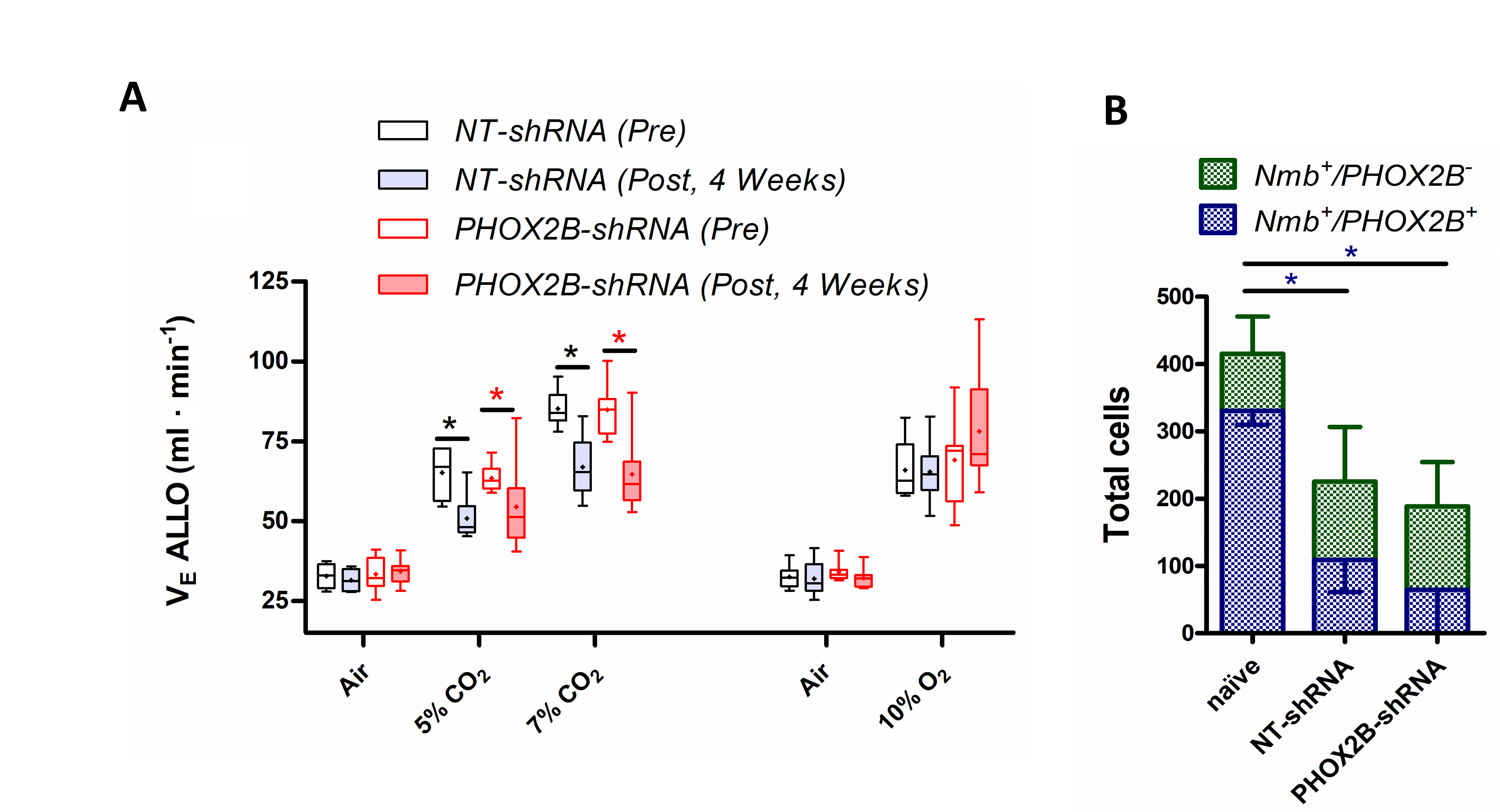
