## supplemental table 1 for "Knockdown of PHOX2B in the retrotrapezoid nucleus reduces the central CO_2_ chemoreflex in rats"

| Figure Reference | Conditions Compared | Main ANOVA | Test |
| --- | --- | --- | --- |
| Fig. 1A - FR | Room Air | Pre vs. Postlesion: F(1, 39) = 11.591, p = 0.002 | Mixed ANOVA |
|  | 5% CO_2_ | Pre vs. Postlesion: F(1, 39) = 32.623, p < 0.001 |  |
|  | 7.2% CO_2_ | Pre vs. Postlesion: F(1, 39) = 16.320, p < 0.001 |  |
| Fig. 1B - VT | Room Air | NS | Mixed ANOVA |
|  | 5% CO_2_ | NS |  |
|  | 7.2% CO_2_ | Interaction: F(1, 39) = 5.523, p < 0.008 |  |
| Fig. 1C – VE ALLO | Room Air | Pre vs. Postlesion: F(1, 39) = 4.962, p < 0.032  Interaction: F(1, 39) = 5.288, p < 0.009 | Mixed ANOVA |
|  | 5% CO_2_ | NS |  |
|  | 7.2% CO_2_ | NS |  |
| Fig. 1D VO2 | Room Air | NS | Mixed ANOVA |
|  | 5% CO_2_ | NS |  |
|  | 7.2% CO_2_ | NS |  |
| Fig. 1E – VE VO2 ALLO | Room Air | NS | Mixed ANOVA |
|  | 5% CO_2_ | NS |  |
|  | 7.2% CO_2_ | NS |  |
| Fig. 1F - HCVR | 5% CO_2_ | NS | Mixed ANOVA |
|  | 7.2% CO_2_ | NS |  |
| Fig. 2C | Total cells | F(2,11) = 145.0, p < 0.001 | One Way ANOVA |
|  | Nmb+/PHOX2B+ | F(2,11) = 85.47, p < 0.001 |  |
|  | Nmb+/PHOX2B- | NS |  |
| Fig. 3C | Total cells | F(2,19) = 6.356 p = 0.0087 | One Way ANOVA |
|  | Nmb+/PHOX2B+ | F(2,19) = 28.24, p < 0.001 |  |
|  | Nmb+/PHOX2B- | F(2,19) = 78.18, p < 0.001 |  |
| Fig. 4A - FR | Room Air | Pre vs. Postlesion: F(1, 21) = 14.742, p = 0.001 | Mixed ANOVA |
|  | 5% CO_2_ | Pre vs. Postlesion: F(1, 21) = 21.000, p < 0.001 |  |
|  | 7.2% CO_2_ | Pre vs. Postlesion: F(1, 21) = 29.221, p < 0.001 |  |
| Fig. 4B - VT | Room Air | Pre vs. Postlesion: F(1, 21) = 12.510, p = 0.002  Interaction: F(1, 21) = 6.462, p = 0.007 | Mixed ANOVA |
|  | 5% CO_2_ | Interaction: F(1, 21) = 3.305, p = 0.057 **(NS)*** |  |
|  | 7.2% CO_2_ | Pre vs. Postlesion: F(1, 21) = 8.926, p = 0.007  Interaction: F(1, 21) = 9.776, p = 0.001 |  |
| Fig. 4C – VE ALLO | Room Air | NS | Mixed ANOVA |
|  | 5% CO_2_ | Pre vs. Postlesion: F(1, 21) = 4.159, p = 0.054 **(NS)*** |  |
|  | 7.2% CO_2_ | Interaction: F(1, 21) = 6.928, p = 0.005 |  |
| Fig. 4D VE VO2 ALLO | Room Air | Interaction: F(1, 21) = 3.643, p = 0.044 | Mixed ANOVA |
|  | 5% CO_2_ | Treatment: F(1, 21) = 4.459, p = 0.024 |  |
|  | 7.2% CO_2_ | Interaction: F(1, 21) = 3.544, p = 0.047 |  |
| Fig. 4E – HCVR | 5% CO_2_ | NS | Mixed ANOVA |
|  | 7.2% CO_2_ | Interaction: F(1, 21) = 7.737, p = 0.003 |  |
| Fig. 4F – HCVR 7.2% | 7.2% CO_2_ | Naïve: (NS)  NT-shRNA: (NS)  PHOX2B-shRNA: F(2.17) = 6.9, p = 0.0075 |  |
| Fig. 5B | TH+/PHOX2B+ | NS | Repeated Measures  ANOVA |
| Fig. 5D | Nmb CTCF ratio | NS | Repeated Measures  ANOVA |
| Fig. 5E - HCVR 7.2% | Nmb+/PHOX2B+ | R^2^=.739, F(1,19) = 54.670, p < 0.001; Cells (β) = 0.125, p <0.001 | Linear regression |
| Fig. 5F - HCVR 7.2% | Nmb+/PHOX2B- | R^2^=.482, F(1,19) = 18.649, p < 0.001; Cells (β) = -0.185, p <0.001 | Linear regression |
| Fig. 6A | Gpr4 CTCF | F(2,11) = 48.41, p = 0.002 | Repeated Measures  ANOVA |
| Fig. 6B | Gpr4 CTCF | F(7,47) = 2.729, p = 0.0228 | Repeated Measures  ANOVA |
| Fig. 6E | Task2 CTCF | F(1,11) = 17.05, p = 0.0034 | Repeated Measures  ANOVA |
| Fig. 6F | Task2 CTCF | F(7,47) = 2.598, p = 0.0287 | Repeated Measures  ANOVA |
| Fig. S1A | Room Air | NS | Mixed ANOVA |
|  | 5% CO_2_ | Pre vs. Postlesion: F(1, 12) = 23.460, p < 0.001 |  |
|  | 7% CO_2_ | Pre vs. Postlesion: F(1, 12) = 43.570, p < 0.001 |  |
|  | Room Air | NS |  |
|  | 10% O_2_ | NS |  |
| Fig. S1B | Nmb+/PHOX2B+ | F(2,11) = 17.642, p = 0.001 | One Way ANOVA |
|  | Nmb+/PHOX2B- | NS |  |

**Table S1. Details on the main statistical analyses.**
